# Social isolation-induced epigenetic and transcriptional changes in *Drosophila* dopaminergic neurons

**DOI:** 10.1101/443226

**Authors:** Pavan Agarwal, Phuong Chung, Ulrike Heberlein, Clement Kent

## Abstract

Epigenetic mechanisms play fundamental roles in brain function and behavior and stressors such as social isolation can alter animal behavior via epigenetic mechanisms. However, due to cellular heterogeneity, identifying cell-type-specific epigenetic changes in the brain is challenging. Here we report first use of a modified INTACT method in behavioral epigenetics of *Drosophila*: a method we call mini-INTACT. Using ChIP-seq on mini-INTACT purified dopaminergic nuclei, we identified epigenetic signatures in socially-isolated and socially-enriched *Drosophila* males. Social experience altered the epigenetic landscape in clusters of genes involved in transcription and neural function. Some of these alterations were predicted by expression changes of four transcription factors and the prevalence of their binding sites in several clusters. These transcription factors were previously identified as activity-regulated genes and their knockdown in dopaminergic neurons reduced the effects of social experience on sleep. Our work enables the use of *Drosophila* as a model for cell-type-specific behavioral epigenetics.

## Introduction

Environmental stressors have robust effects on the behavior of animals including humans, rodents and fruit flies. Social isolation is considered a form of ‘passive’ stress that can profoundly affect behaviors by inducing anxiety and depression-like symptoms (Grippo et al., 2007; Hall, 1998; Wallace et al., 2009). For instance, solitary confinement in humans has been shown to induce depressive symptoms, increased aggression (Ferguson et al., 2005) and increased risk for suicide (Kaba et al., 2014; Reeves and Tamburello, 2014). In addition, social isolation is known to affect sleep quality and duration in humans (Cacioppo et al., 2000; Friedman, 2011), mice (Febinger et al., 2014; Greco et al., 1988) and the fruit fly *Drosophila melanogaster* (Brown et al., 2017; Ganguly-Fitzgerald et al., 2006).

Epigenetic mechanisms engaged by stressors such as early life adversity (Champagne, 2010; McGowan and Szyf, 2010), reduced maternal care (Weaver et al., 2004), maternal separation (Pusalkar et al., 2015; Sasagawa et al., 2017), drugs of abuse (Chase and Sharma, 2013; Gozen et al., 2013; Jung et al., 2016; Renthal et al., 2009; Wang et al., 2007), and social defeat (Valzania et al., 2017) play a key role in influencing gene expression in the brain. Social isolation has been shown to cause epigenetic changes in the midbrain of mice (Siuda et al., 2014) and an increase in DNA methylation in dopaminergic neurons (Niwa et al., 2013; Niwa et al., 2016). Several studies have implicated dopaminergic neurons in the effects of social isolation in rodents (Hall et al., 1998; Jones et al., 1992; Sasagawa et al., 2017) and social isolation has been shown to decrease dopamine levels in flies (Ganguly-Fitzgerald et al., 2006) and mice (Niwa et al., 2013). Dopaminergic neurons play an important role in modulating behaviors influenced by social isolation in *Drosophila*, including aggression (Alekseyenko et al., 2013), sleep (Ganguly-Fitzgerald et al., 2006; Liu et al., 2012a; Pimentel et al., 2016; Sitaraman et al., 2015; Ueno et al., 2012), and alcohol intoxication (Bainton et al., 2000). It is not known, however, how stressors such as social isolation influence the epigenome in specific cell types of the brain to affect behavior.

The brain is a highly heterogeneous tissue. This poses a challenge for epigenomic studies since ChIP-seq and RNA-seq data obtained from brain tissue are significantly more variable than data obtained from other tissue types or cells in culture (Maze et al., 2014). This is especially challenging for small model organisms such as *Drosophila*, where manually dissecting subsets of brain regions for epigenomic analysis is not possible. Consequently, studies of behavioral epigenetics in *Drosophila* have used either mutants or flies in which the GAL4-UAS system (Brand and Perrimon, 1993) was used to modulate levels of epigenetic writers or erasers (Anreiter et al., 2017; Anreiter et al., 2019; Fitzsimons et al., 2013; Gupta et al., 2017; Johnson et al., 2010; Koemans et al., 2017; Kramer et al., 2011; Perry et al., 2017; Schwartz et al., 2016; Taniguchi and Moore, 2014; van der Voet et al., 2014; Xu et al., 2014b). Studies that looked at global epigenetic changes using ChIP-seq have used either entire fly heads or whole animals after drug treatment or epigenetic mutation (Ghezzi et al., 2013; Kramer et al., 2011; Wang et al., 2007).

Strategies to isolate specific cell types from brains, such as laser capture microdissection (Emmert-Buck et al., 1996) or manual sorting of neurons (Hempel et al., 2007; Nagoshi et al., 2010) do not provide enough material for epigenomic analysis. A popular approach for cell type-specific epigenomic analysis is INTACT (isolation of <u>n</u>uclei <u>ta</u>gged in specific <u>c</u>ell types) (Deal and Henikoff, 2010). INTACT allows the isolation of specific cell types using tagged nuclei that are affinity purified from a heterogeneous cell population. Recent advances in INTACT have made it possible to use this method in *C. elegans* (Steiner et al., 2012), *Drosophila* (Henry et al., 2012), and mouse (Mo et al., 2015; Mo et al., 2016) for epigenomic and proteomic (Amin et al., 2014) analyses. Despite its versatility, to the best of our knowledge, no studies to date have utilized INTACT for analysis of rare cell types in the field of behavioral epigenetics. INTACT in mouse has been shown to work with 1-3% of total adult neuronal nuclei (Mo et al., 2015) and epigenetic analysis with ChIP-seq required ~0.5-1 million purified neuronal nuclei (Mo et al., 2016). INTACT in *Drosophila* either requires thousands of animals to access rare cell types (Henry et al., 2012) or the use of pan-neuronal or pan-glial drivers to obtain sufficient nuclei for epigenetic analysis (Henry et al., 2012; Ma and Weake, 2014; Ye et al., 2017). This represents a significant barrier for the field of behavioral epigenetics, in which rare cell types need to be collected in restricted time windows and where collecting tissue from large number of animals would be difficult.

To address these issues, we developed a modification of the INTACT method, mini-INTACT, which uses approximately 100-fold less material. We used mini-INTACT to purify nuclei from dopaminergic neurons, which comprise less than 0.1% of fly brain neurons. We used 200-250 fly heads (10-15,000 nuclei) of socially-isolated and socially-enriched flies and ascertained epigenetic changes on a genome-wide scale using ChIP-seq. Comparing the enrichment profiles of six different histone modification marks with mRNA expression levels in dopaminergic neurons obtained by RNA-seq revealed clusters of genes that may contribute to the effects of social isolation and social enrichment. Our unsupervised clustering analysis followed by gene ontology (GO) analysis of these groups showed an enrichment of genes encoding readers and writers of the epigenome, cell signaling molecules, and molecules involved in neural and behavioral processes. We found that some genes encoding activity-regulated transcription factors (ARG-TFs) (Chen et al., 2016) respond to social environment in dopaminergic neurons, and that knockdown of the genes encoding four of these ARG-TFs (*cabut, Hr38, stripe, CrebA*) reduced the effects of social experience on daytime sleep. Taken together, these data show that the epigenetic landscape of dopaminergic neurons undergoes modifications with just four days of social isolation in adult male flies and that ARG-TFs are part of these changes.

## Results

### mini-INTACT purifies rare cell types from adult *Drosophila* brain

The INTACT method developed in *Drosophila* expresses a SUN domain protein (UNC84) from *C. elegans* that localizes GFP to the inner nuclear membrane (*unc84-2xGFP*) (Henry et al., 2012). While the INTACT method works well to isolate specific cell types from *Drosophila*, it requires thousands of fly heads to access rare cell types. This represents a significant challenge for the field of behavioral epigenetics, where animals need to be perturbed and collected in restricted temporal windows, and where manually manipulating large number of animals is difficult. To address this issue, we modified the INTACT method to isolate rare cell types (<0.1% of adult *Drosophila* brain) from 200-250 fly heads; we named this modified method mini-INTACT (Figure 1A and Materials and Methods).

**Figure 1:**
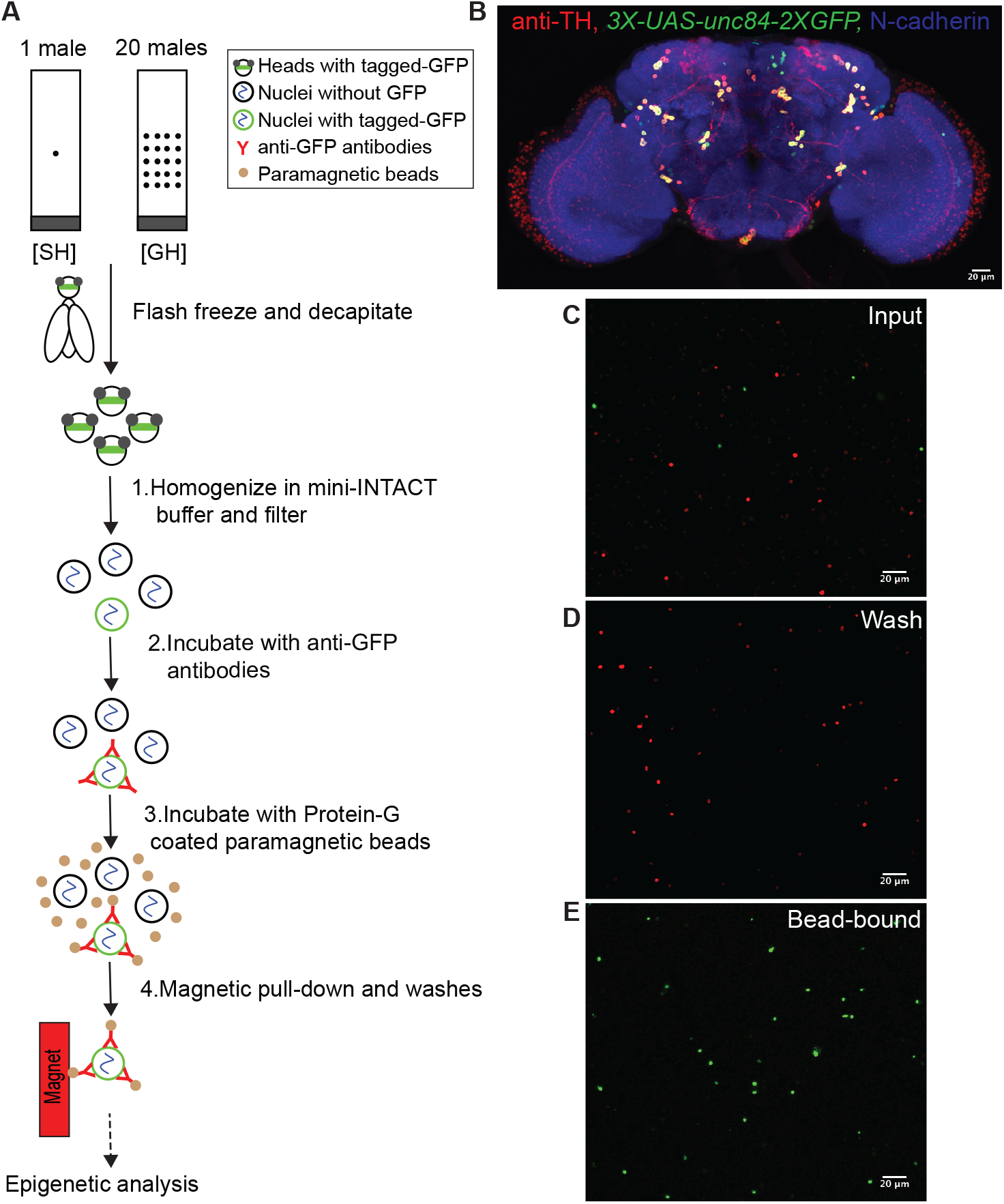
mini-INTACT method affinity purifies cell-type specific nuclei for epigenetic analysis in *Drosophila.* (A) Schematic of mini-INTACT method to affinity purify dopaminergic neurons from heads of *Drosophila* male flies after four days of social isolation or social enrichment. (B) Expression of SUN-tagged GFP (*3XUAS-unc84-2XGFP*) in dopaminergic neurons driven by *TH-GAL4* in an adult male brain. (C) To assess the purity of dopaminergic nuclei, nuclei were obtained from a mixture of heads derived from flies expressing *10XUAS-unc84-tdTomfl* under the control of the pan-neuronal *elav-GAL4* driver (red) and flies expressing *3XUAS-unc84-2XGFP* under the control of the *TH-GAL4* driver (green). (D) After capturing green nuclei using bead-bound anti-GFP antibodies, red nuclei were washed away. (E) Bead-bound green nuclei were almost devoid of contaminating red nuclei.

To achieve this ~50-100-fold reduction in input material, we made several changes to the protocol (see Materials and Methods for details), including an improved homogenizer design to prevent sample loss (Figure 1- Figure Supplement 1); a 20-fold reduction in homogenization and immunoprecipitation volume; the use of a single buffer system for homogenization, immunoprecipitation and washing; and the sequential addition of anti-GFP antibodies and magnetic beads directly to the homogenate for increased binding efficiency.

We expressed the INTACT transgene in dopaminergic neurons using the tyrosine hydroxylase driver, *TH-GAL4* (Friggi-Grelin et al., 2003), which is expressed in ~120 neurons in the adult brain (Azanchi et al., 2013; Friggi-Grelin et al., 2003; White et al., 2010)(Figure 1B). We compared expression of the *TH-GAL4*-driven transgene *UAS-UNC84-2XGFP* in the adult brain after varying the copy number of the upstream activator sequences (UAS) from 3X to 5X and 10X. The *3X-UAS-unc84-2XGFP* transgene most faithfully reproduced *TH*-*GAL4* expression (Figure 1B); ectopic expression was seen when 5 or 10 copies of UAS-tagged GFP were used (Figure 1-Figure Supplement 2). Therefore, we used *3X-UAS-unc84-2XGFP* for all our experiments. Social isolation affects daytime sleep (Brown et al., 2017; Ganguly-Fitzgerald et al., 2006); we therefore tested for the effects of expression of the INTACT transgenes on daytime sleep using the *Drosophila* activity monitor. Expression of *3X-UAS-unc84-2XGFP* in dopaminergic neurons did not affect daytime sleep (Figure 1-Figure Supplement 3), leading us to conclude that the expression of the transgenes had no significant effects on fly behavior.

To assess purity of the isolated nuclei, we mixed 200 heads of flies expressing *UAS-UNC84-2XGFP* driven by *TH-GAL4* with 200 heads of flies expressing *UAS-UNC84-tdTomfl* driven by the pan-neuronal driver *elav*-*GAL4*. Processing these heads using mini-INTACT resulted in a ratio of ~120 GFP-positive green to 10^5^ tdTomfl-positive red nuclei. Very few red nuclei were observed in the purified bead-bound sample as compared to green nuclei (Figure 1C-E). Therefore, the purity obtained by mini-INTACT (~98%, Figure 1- Figure Supplement 4) is comparable to that described for the INTACT method (Henry et al., 2012) that requires ~50-100 times more input material.

By manually counting various purified and diluted samples we assessed the yield of nuclei to be in the range of 30-50% (data not shown). Therefore, from the heads of 200-250 flies, we estimated a yield of 10,000-15,000 dopaminergic nuclei for each ChIP-seq reaction. Dopaminergic neurons were obtained from *Drosophila* males that were either socially isolated or socially enriched for four days, hereafter referred to as single-housed (SH) and group-housed (GH) male flies, respectively. Chromatin was processed from these nuclei for ChIP-seq using six different histone modification marks as described in the Materials and Methods and below.

In summary, mini-INTACT allowed us to retrieve sufficient chromatin for ChIP-seq analysis of six histone marks from dopaminergic neurons of 200-250 flies for each housing condition.

### Epigenomic profiling of dopaminergic neurons from socially isolated and socially enriched male flies

The genome-wide profiles of activating and repressive marks (David et al., 2015) with respect to gene bodies are shown in ngs.plot displays (Shen et al., 2014) averaged over the genome (Figure 2A-F). As expected from previous studies with human cells (Barski et al., 2007; Mikkelsen et al., 2007), flies (Kharchenko et al., 2011; Kramer et al., 2011), and mouse brain (Feng et al., 2014), activating marks H3K4me3, H3K27ac and H3K9/K14ac were maximally enriched downstream of the transcription start site (TSS) (Figure 2A-C), while H3K36me3, which has been associated with transcriptional elongation, is skewed towards transcription end site (TES) with enrichment in the gene body (Figure 2D). Repressive marks H3K9me3 and H3K27me3 were depleted from TSS and TES and enriched in the central portion of the gene body (Figure 2E-F).

**Figure 2:**
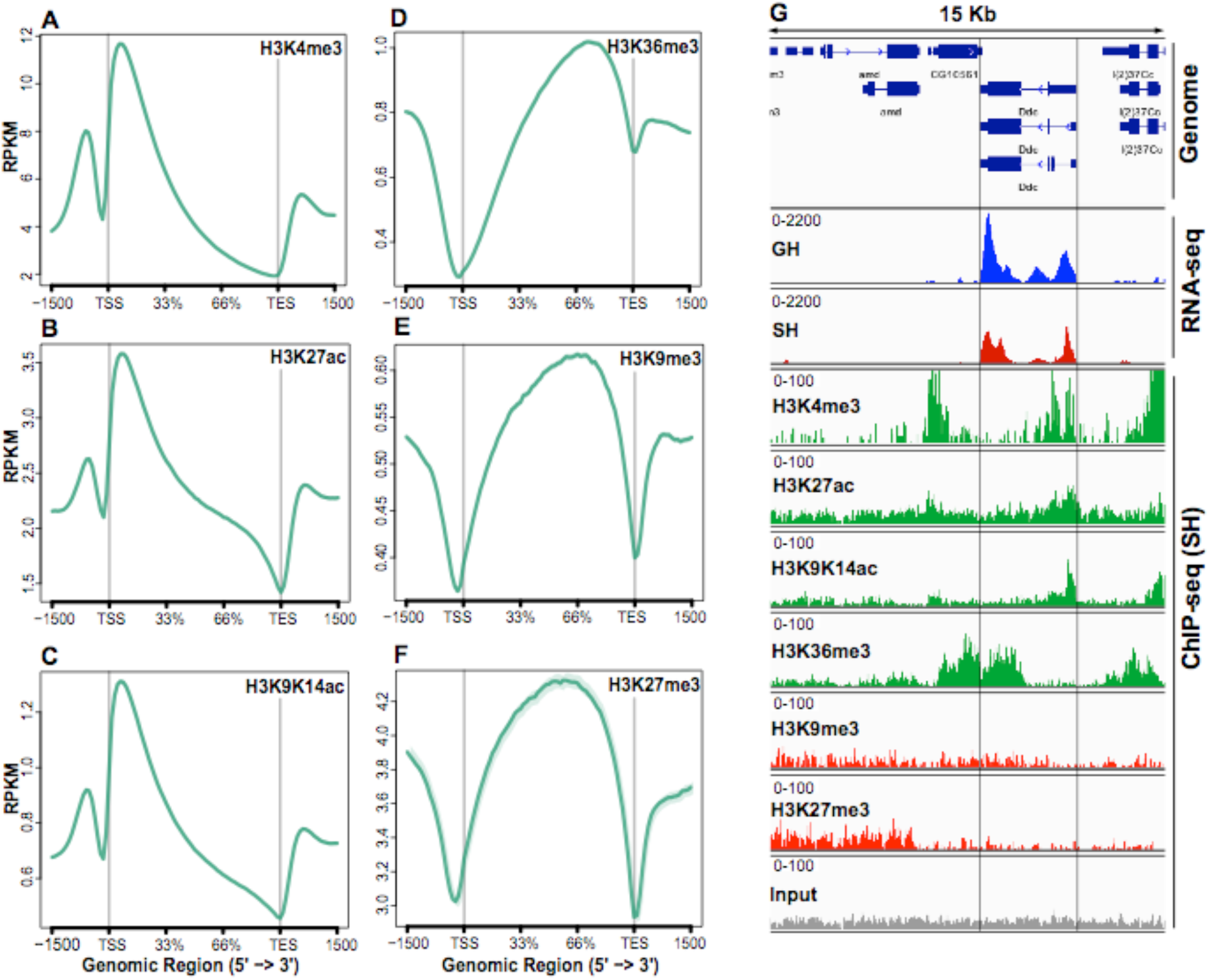
Epigenome of mini-INTACT purified dopaminergic neurons measured by ChIP-seq and RNA-seq. (A-F) Genome-wide profiles of the levels for the six epigenetic marks shown as ngs.plot displays. (A-C) Activating marks were concentrated in the promoter and immediately downstream of the TSS. (D) H3K36me3, a mark associated with transcriptional elongation, was enriched in the gene body and skewed towards the TES. (E-F) Two repressive marks were depleted from the TSS and TES, concentrated in the gene body, and enriched upstream of the promoter region. (G) Epigenetic and transcriptional enrichment profiles surrounding the *Ddc* gene. The RNA-seq panels show that *Ddc* is more strongly expressed in GH (blue) than in SH (red) males. The distribution of epigenetic marks shown are representative of the SH dataset. The four activating marks (green panels) were high in this strongly expressed gene, while the two repressive marks (red) showed low levels. An example of single “input” DNA track, which is used as a control for mark levels, is shown in the final panel.

As an example for transcriptional and epigenetic changes at a specific locus, we depict the highly-expressed *Dopa decarboxylase* (*Ddc*) gene, which is involved in dopamine synthesis. *Ddc* mRNA levels were upregulated in GH flies as compared to SH flies (Figure 2G), which is consistent with a previous study showing that the levels of dopamine are lower in the heads of socially-isolated flies (Ganguly-Fitzgerald et al., 2006). The epigenetic profile of this locus recapitulates the global profile, with marks associated with transcriptional activation (H3K4me3 and H3K27ac) centered around the TSS, H3K36me3 skewed towards the TES, and repressive marks H3K9me3 and H3K27me3 not showing enrichment as compared to input DNA. Comparative analysis of epigenetic profiles between GH and SH males using SICER (Zang et al., 2009) showed that the levels of the activating mark H3K4me3 were significantly higher in GH flies around the *Ddc* gene (normalized read count GH: 35.10, SH:30.58, p=0.0002, p_adjusted_=0.0004), and that the activating mark H3K27ac was similarly increased (GH: 1069, SH: 651, p=1.5×10^-107^, p_adjusted_<10^-60^) in agreement with the pattern of mRNA expression. Repressive marks, which were already very low on this gene, showed no significant differences.

ChIP-seq replicates for histone modification marks were highly correlated (median Pearson’s r of log-transformed coverage among all pairs of biological replicates, r>0.99 (Figure 2- Figure Supplement 1). The genome-wide correlation between levels of activating and repressive marks with each other and with mRNA levels is shown in Table 1. All activating mark levels correlate positively with each other and with mRNA levels, while repressive marks correlate positively with each other and negatively with mRNA levels, as expected. H3K9me2 and H3K9me3 modifications are associated with Heterochromatin Protein 1 (HP1)-mediated heterochromatin formation and transcriptional repression (David et al., 2015), however these modifications are not strongly correlated with transcriptional repression in either human cells (Barski et al., 2007) or *Drosophila* (Kramer et al., 2011). Consistent with these findings, we find correlations of H3K9me3 to be weaker with mRNA levels and with activating marks when compared with the repressive mark H3K27me3.

**Table 1:** Pearson correlation coefficient values of pairwise comparisons among ChIP-seq for six histone modification marks and gene expression. Activating epigenetic marks and positive correlations are shown in green, repressive marks and negative correlation are shown in red.

Tables and Figures with legends
|  | H3K27ac | H3K9ac_K14ac | H3K36me3 | H3K9me3 | H3K27me3 | mRNA |
| --- | --- | --- | --- | --- | --- | --- |
| H3K4me3 | 0.76 | 0.84 | 0.73 | -0.46 | -0.55 | 0.54 |
| H3K27ac |  | 0.85 | 0.62 | -0.29 | -0.34 | 0.41 |
| H3K9ac_K14ac |  |  | 0.61 | -0.25 | -0.33 | 0.47 |
| H3K36me3 |  |  |  | -0.35 | -0.42 | 0.38 |
| H3K9me3 |  |  |  |  | 0.90 | -0.32 |
| H3K27me3 |  |  |  |  |  | -0.38 |

Analysis of ChIP-seq data using SICER (Zang et al., 2009) returned thousands of “islands” in which epigenetic mark levels were significantly different between GH and SH males (FDR<0.001) (Supplement Table 1). Typically, an island does not cover the entirety of a gene, so interpretation of SICER islands requires care. For example, an H3K4me3 island with a fold change of 1.25 was found within the body of the *foraging* gene, from position 3,622,074 to 3,656,953 bp on chromosome 2L. This island covers the first exon of seven *foraging* transcripts, but not of the remaining 6 transcripts annotated in Flybase (Dos Santos et al., 2015)(www.flybase.org). By contrast, an island in *Snmp2* covers half of the first exon of all three transcripts and has H3K4me3 fold change of 1.70. Details of SICER-detected islands are in Supplement Table 1.

In summary, when averaged over entire gene bodies there are small but statistically significant changes in histone marks, but when examined in islands detected by SICER there are much larger changes, often restricted to regions such as the first exon of a gene (for activating mark H3K4me3).

### Social experience induces transcriptional changes in dopaminergic neurons

Since most of the transcripts are exported from the nucleus soon after transcription (Rodriguez et al., 2004), nuclear RNA alone may not represent the transcriptional changes due to a four-day-long social experience. A recent study showed that considerable differences exist in the profiles of nuclear and cytosolic transcripts of individual cells (Abdelmoez et al., 2018). Therefore, to profile both nuclear and cytosolic mRNAs, we isolated dopaminergic neurons from GH and SH males using fluorescence activated cell sorting (FACS) and performed RNA-seq (see Materials and Methods). Replicate concordance was assessed using Pearson’s r of log-transformed counts among all pairs of replicates (r=0.95 for GH and r=0.91 for SH flies). These correlations are similar to those reported before for RNA-seq from dopaminergic neurons (Abruzzi et al., 2017; Chen et al., 2016).

We used three methods (EdgeR, CyberT, and FCros) to identify genes that are differentially expressed in dopaminergic neurons of GH and SH flies (Dembélé and Kastner, 2014; Kayala and Baldi, 2012; Robinson et al., 2010). EdgeR and CyberT use generalizations of the between-treatment t-test method, while FCros uses a nonparametric method based on fold changes, which is more robust to variation in mRNA counts. Using EdgeR with a FDR of 5% (see Materials and Methods), we identified 16 genes upregulated in SH and 9 genes upregulated in GH males (Supplement Table 2). The fold-change based technique FCros identified 451 genes upregulated in SH and 466 upregulated in GH, after FDR correction of 5%. CyberT produced intermediate results. In figures and tables, we quote the FDR 0.05 obtained with FCros values, except where noted otherwise. Supplement Table 2 shows each gene reported as differentially expressed by any of the 3 methods.

Gene Ontology analysis of all differentially-expressed genes (upregulated in both GH and SH males) using the DAVID GO tool (Huang et al., 2009) found two related GO groups: epigenetic (unadjusted p=0.0098) and negative regulation of gene expression (p=0.016). GOrilla GO analysis (Eden et al., 2009) found peptide n-acetyltransferase activity (p=0.00044), the latter group containing genes belonging to several histone acetyltransferase complexes genes including Tip60 complex members Enhancer of Polycomb and *dom*, SAGA complex members *Taf10b* and *Taf12*, TAC1 complex members *nejire* and *Sbf*, and Enok complex members *enok*, *Gas41* and *Ing5*. The full GOrilla analysis is shown in Supplement Data 1.

Daytime activity is significantly higher in SH flies as compared to GH flies (Ganguly-Fitzgerald et al., 2006), suggesting that metabolic activity might be higher in SH flies. It is also known that mitochondrially-encoded genes are upregulated in waking flies (Cirelli and Tononi, 1998). Consistent with these observations, in our RNA-seq dataset we found that of 15 known mitochondrially encoded genes, 14 were higher in SH than in GH flies (p=0.0005, binomial test).

In summary, transcript levels of many genes expressed in dopaminergic neurons were changed by social housing conditions, including those of many epigenetic reader and writer genes.

### Social experience alters epigenetic landscape

To understand how social experience affects the epigenetic landscape of dopaminergic neurons, we focused on epigenetic changes observed in the top 40% of genes by mRNA expression levels (“expressed genes”). We clustered the z-score normalized differences between GH and SH flies for all 6 epigenetic marks and for mRNA, and performed k-means clustering as in Shen *et al.* (2013)(see Materials and Methods and Figure 3- Figure Supplement 1). Eight clusters provided optimal separation of genes (elbow test). These clusters, arranged in increasing order of mean mRNA expression levels, are shown as a heat map of mRNA and epigenetic mark z-score values in Figure 3, with red showing marks/mRNAs that are higher in GH flies and blue showing those that are higher in SH flies.

**Figure 3:**
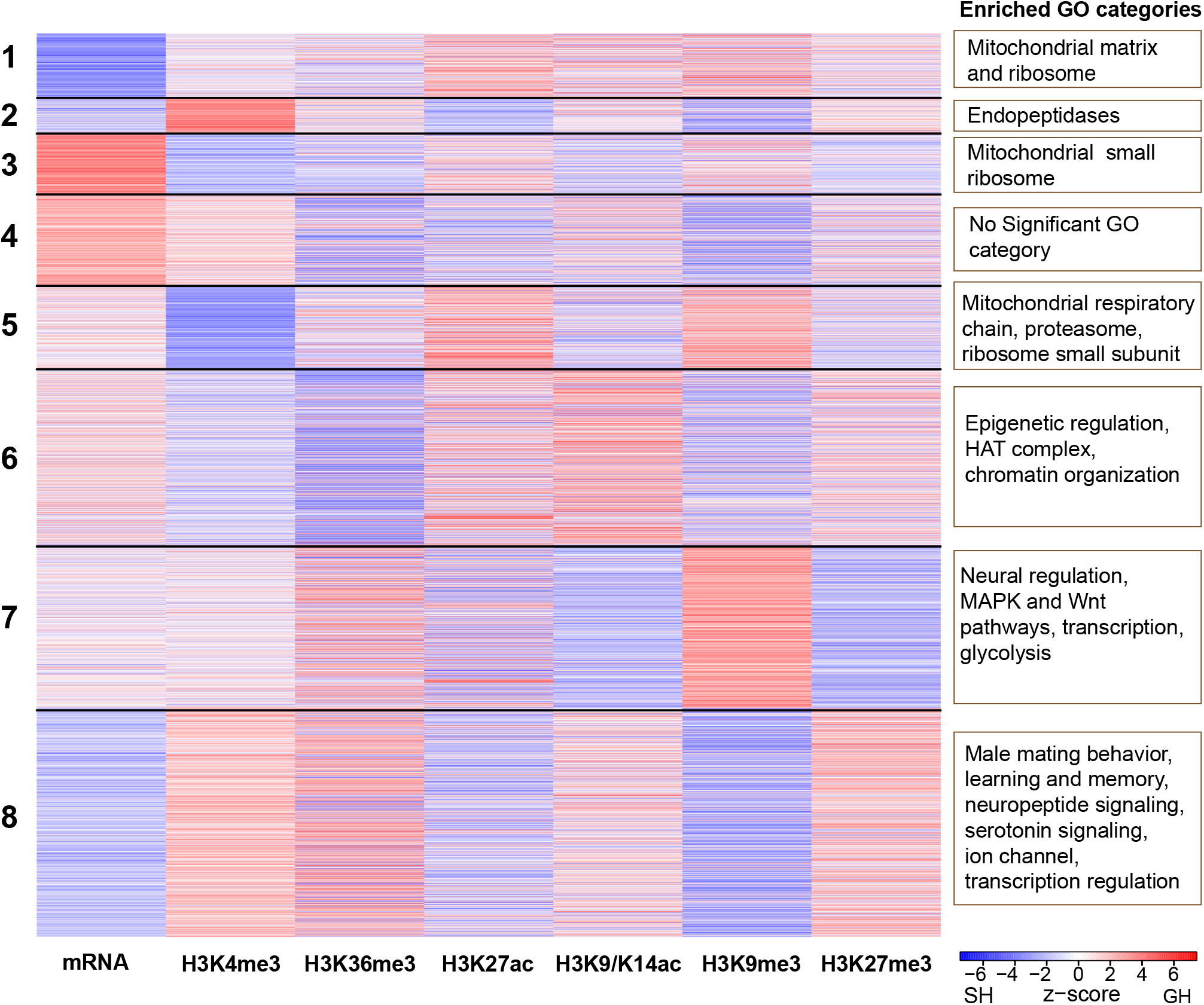
Epigenetic landscape of genes expressed in dopaminergic neurons is modulated by social experience. Heat map of eight groups identified by k-means clustering of the change in mark and mRNA levels between GH and SH males. Red lines show genes whose marks or mRNA was higher in GH than SH males, blue lines show those that were higher in SH than GH males. Some clusters are enriched for genes with neural and regulatory functions, especially clusters 6-8. Enriched GO categories from each cluster are shown on the right.

The first five clusters are enriched for house-keeping functions, and include mitochondrial, ribosomal, and proteasome genes. However, the last three clusters (6-8, containing genes with higher expression) are enriched in neural and regulatory functions.

Cluster 6 is enriched for genes with epigenetic functions, including histone acetyltransferase (HAT) genes. As noted above, HAT genes and several epigenetic regulators encode differentially expressed mRNAs; but as can be seen in the left-hand column (mRNA z-score), mRNA level changes are heterogeneous, as some genes in this cluster are upregulated in GH males (red) and others in SH males (blue). This is interesting considering that the k-means analysis grouped genes in this cluster not primarily by the direction of their mRNA change with regard to housing condition, but by their epigenetic mark changes; this cluster is enriched for readers and writers of epigenetic marks.

The seventh cluster is enriched for genes regulating neural function (some of which are members of the MAPK or WNT signaling pathways), transcription factors, and glycolysis genes. In this cluster there is a pair of marks that show strong, anti-correlated changes: Heterochromatin Protein 1 (HP1)-associated H3K9me3 (higher in GH than SH) and the Polycomb repressive complex 2 (PRC2)-associated H3K27me3 mark (higher in SH than GH).

The two inhibitory marks also change in opposite directions in the final (highest expression) cluster 8, but in this cluster the directions of change are reversed. H3K9me3 in cluster 8 is higher in SH than GH males and H3K27me3 is higher in GH than SH males. This cluster is enriched in neural function genes, including those involved in male mating behavior, learning and memory, synaptic, neuropeptide and serotonin signaling, as well as ion channels and transcription regulation genes. Genes of this cluster have on average higher expression in SH flies than in GH (p<10^-15^, t=-15.06, df=1361).

In summary, there are clusters of genes whose epigenetic marks and mRNA levels respond to social experience in similar ways within each cluster, but quite differently between clusters. This suggests that different regulatory programs may be acting in each cluster. We use this putative division of genes into epigenetically distinct clusters to try to determine what the regulatory program might be in the next section.

### Social enrichment induces activity-regulated genes in dopaminergic neurons

We used the Centrimo tool (Bailey and Machanick, 2012) to search for transcription factors (TFs) whose binding sites might be enriched (occur more often than chance) in promoter-proximal regions of genes expressed in dopaminergic neurons. The 8 epigenetic clusters discussed in the previous section provided us with groups of genes that had similar regulatory programs (as evidenced by their epigenetic and transcriptional response to housing conditions). We used Centrimo to search for TFs whose binding sites were enriched in genes of each cluster relative to a control group of the same number of genes randomly chosen from other clusters. The promoter-proximal region (±500 bp from TSS) was scanned. We found a group of 24 TF motifs that were enriched in one or more of the clusters’ promoter regions. These correspond to 14 TF genes, as in many cases multiple binding motifs are documented for one TF in the motif databases used by Centrimo (Supplement Data 2).

We further filtered the TFs under investigation by two criteria: 1) the TF had to be in the expressed gene set and 2) the TF had to show at least a 33.3% change in transcript levels in response to housing conditions. Five TFs met our criteria: Hr38 (Hormone receptor-like in 38), Sr (Stripe), CrebA, Cbt (Cabut), and Pho (Pleiohomeotic). Interestingly, the genes encoding four of these TFs (*Hr38*, *sr*, *CrebA* and *cbt*) are orthologs of vertebrate immediate early genes (Hu et al., 2011). The expression of these genes was higher in GH males than in SH males. We hypothesized, consistent with another study (Ganguly-Fitzgerald et al., 2006), that being in the GH environment constitutes an enrichment of stimuli for male flies. In a recent study (Chen et al., 2016) the Rosbash group thermogenetically stimulated dopaminergic neurons by expressing dTRPA1 using the *TH-GAL4* driver and measured changes in mRNA levels after 60 minutes. Genes with large transcriptional upregulation due to neuronal stimulation were called “Activity Related Genes” (ARGs). We compared the change in expression levels (log-fold change) of the top 50 upregulated ARGs in dopaminergic neurons found by Chen *et al.* with the change of gene expression in dopaminergic neurons between GH and SH males in our dataset; we found a significant positive correlation (r=0.41, p=0.003).

Interestingly, changes in the levels of some histone marks observed between GH and SH males also correlated significantly with Chen *et al.’s* changes in expression of ARGs upon neuronal stimulation: H3K9me3 (r=-0.35, p=0.01), H3K27me3 (r=0.46, p=0.0009) and H3K4me3 (r=0.32, p=0.026). This result suggests that genes in dopaminergic neurons responding to short-term direct neural stimulation also respond epigenetically and transcriptionally to the long-term presumed behavioral stimulation of dopaminergic neuron due to interaction among GH flies over the course of four days.

Four transcription factors were among the genes that showed the largest upregulation in response to direct neuronal stimulation: Hr38, sr, CrebA, and Cbt (Chen et al., 2016). All four of these TFs were also upregulated in our RNA-seq data in GH as compared to SH males (Figure 4A), suggesting that they might regulate transcriptional responses of other genes in response to group housing. Interestingly, a recent study of gene expression in the *Drosophila* midbrain found that transcription of these ARGs was correlated across many types of neurons (Croset et al., 2018) under normal conditions – that is, there appears to be a common regulatory program across neural cell types for these genes. The epigenetic effects of social housing on marks were more highly correlated (by 2.4 times) among these ARGs than among all genes (t=2.336, df=20, p=0.03). Of the 10 genes found by Croset *et al.*, 9 were also present in our top 40% expressed genes in dopaminergic neurons (Figure 4, B). These 9 ARGs had GH/SH fold changes ranging from 1.44 to 2.11 (mean 1.70; p=0.004, binomial test; Supplement Table 3). Similarly, genes with log fold change above 1.5 in Figure 4 of Chen *et. al* 2016 had high log fold changes in our data (Figure 4 B, Chen *et al.* high), while lower fold change genes from the same Chen dataset had fold changes in our data not different from zero (Fig. 4 B, Chen *et al.* low) showing that fold change sizes in this set of genes seems to be conserved across experimenters and conditions.

**Figure 4:**
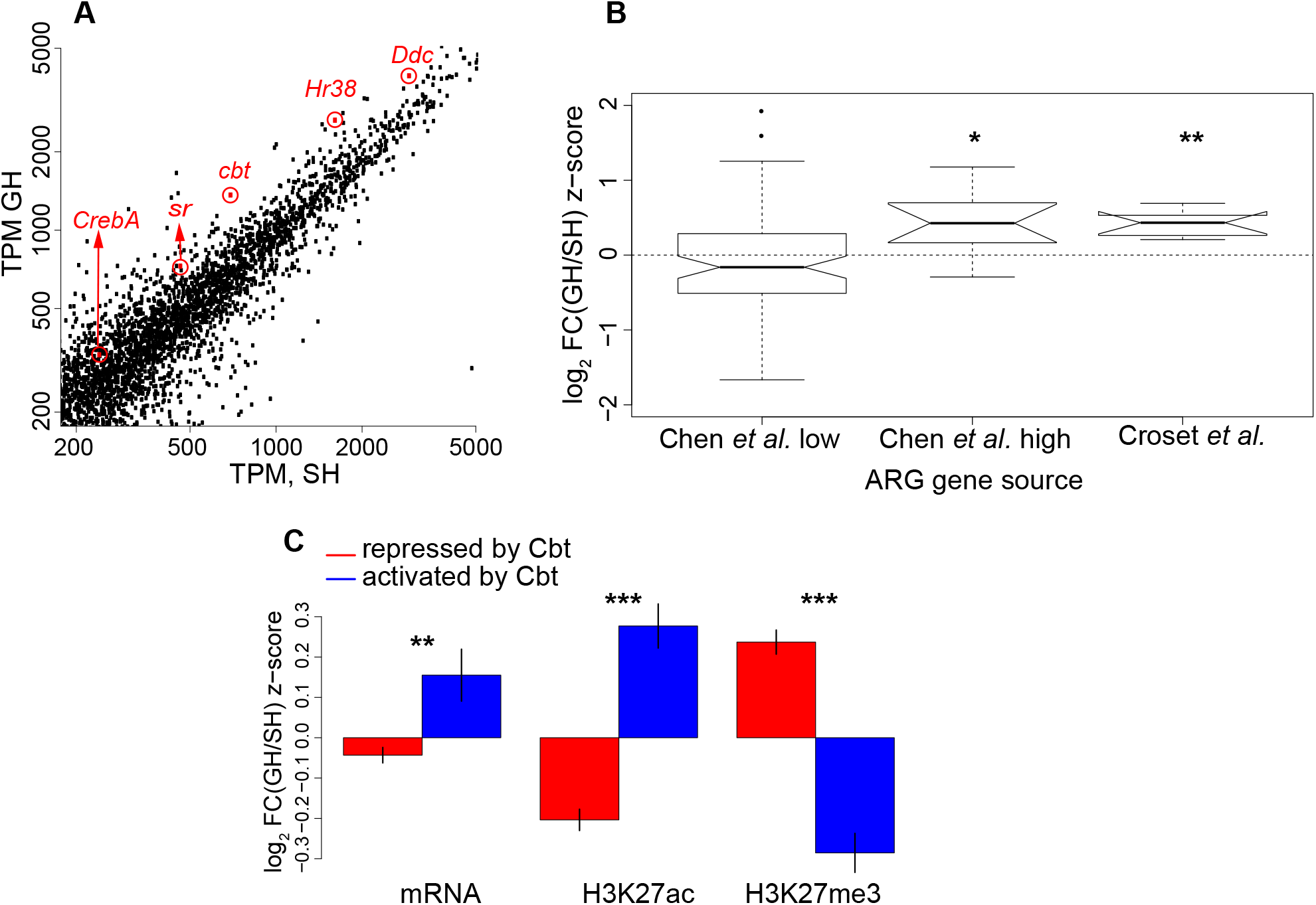
Activity-regulated genes (ARGs) are upregulated in dopaminergic neurons of GH males and correlate with transcriptional repression. (A) A zoomed in scatter plot of GH versus SH mRNA values. *Ddc* (Figure 2) and four ARG-TFs are highlighted. (B) Box plots of mRNA log fold change z-scores (GH is positive, SH is negative) for groups of ARGs from two different studies. Genes with log fold change lower than 1.5 in Chen *et al.* 2016 study are not over-represented in GH flies (Chen *et al.* low). Whereas, the last two groups are significantly over-represented in GH flies. (C) Genes repressed (red) or activated (blue) by the ARG-TF Cbt (from Bartok et al., 2015) are shown on the same z-score scale as in (B). Genes repressed by Cbt have significantly lower mRNA and activating mark H3K27ac, and significantly higher repressive mark H3K27me3. Genes activated by Cbt show the reverse pattern.

In summary, several ARGs expressed in dopaminergic neurons respond similarly to 4 days of social housing and 60 minutes of thermogenetic stimulation. We report below the effects of these ARG transcription factors on downstream targets using both bioinformatic analysis and by manipulating levels of these ARG TFs in dopaminergic neurons and measuring the effect on sleep.

### ARGs predict transcriptional changes due to social experience

It has been suggested that ARGs may in some cases act as transcriptional repressors to fine tune responses to neuronal stimulation (Croset et al., 2018). To test if the factor encoded by the ARG *cbt* is acting as a transcriptional repressor, we compared a published dataset for *cbt* (Bartok et al., 2015) with our data. In the *Bartok et al.* study, genome-wide transcriptional responses were measured upon overexpression and knockdown of *cbt* in adult male fly heads. We used mRNA expression from this study to define two sets of genes: ‘repressed by Cbt’ and ‘activated by Cbt’. The ‘repressed by Cbt’ set contains genes whose expression is increased upon *cbt* knockdown and decreased upon *cbt* overexpression (Supplement Table 4). Conversely, the ‘activated by Cbt’ set contains genes whose expression is decreased upon *cbt* knockdown and increased upon *cbt* overexpression (Supplement Table 5). *cbt* is up-regulated by 94% in dopaminergic neurons of GH males compared to SH males. Hence, if Cbt indeed acts as a transcriptional repressor in dopaminergic neurons, we should see downregulation of genes repressed by Cbt and/or upregulation of genes activated by Cbt in GH males. To test this, we compared gene expression between the two datasets using the top 40% of expressed genes in dopaminergic neurons. Consistent with our hypothesis, we found reduced expression of genes repressed by Cbt in GH males compared to SH males (two-sided Student t: t=-3.31, df=1143, p=0.0001). Genes activated by Cbt were upregulated in GH males, although this effect was not statistically significant (t=1.30, df=269, p=0.196). Thus, Cbt appears to act primarily as a transcriptional repressor in dopaminergic neurons in response to social stimulation: it is higher in GH than in SH neurons, and genes repressed by it in heads (Bartok et al., 2015) are lower in dopaminergic neurons in GH compared to SH males.

We next analyzed the effects of housing on the six histone marks in the two sets of Cbt-regulated genes. For each mark, the difference between the two gene sets was significant at p values ranging from 10^-12^ to 10^-32^. The activating marks H3K4me3, H3K36me3, and H3K9-14ac, and the repressive mark H3K27me3 were higher in GH males in ‘repressed by Cbt’ genes than in ‘activated by Cbt’ genes. By contrast, the marks H3K27ac and H3K9me3 were higher in SH males in the ‘repressed by Cbt’ genes than in the ‘activated by Cbt’ genes (Figure 4C). Interestingly, genes in the ‘repressed by Cbt’ group were over-represented in our eighth k-means cluster (Figure 3) containing genes involved in neuronal function (odds ratio 1.7:1, chi-squared 95.9, df=1, p=1.8*10^-22^). We present a hypothesis for this unusual pattern of mark changes in the Discussion.

If any of the four ARG-TFs (*Hr38, cbt, CrebA* and *sr*) are acting as transcriptional repressors, as suggested by (Croset et al., 2018) and as shown above for *cbt* in dopaminergic neurons, there should be sets of target genes that are down-regulated in GH flies, since these ARG-TFs are up-regulated upon group housing. To test this, we performed multi-linear regressions (see Materials and Methods and Supplement Data 3) with expression level change (mRNA log fold change) between GH and SH males as the dependent variable and the number of TF binding motifs per gene in a 1000 bp region centered on the TSS as the independent variable. The motifs used were for the four TFs in the ARG group and for TF encoded by *pho* (associated with PRC2-mediated epigenetic regulation) (Nitta et al., 2015; Zhu et al., 2011), whose binding motifs were enriched in genes of the 8 clusters described above (Figure 3) (see Materials and Methods and Supplement Data 2).

We did the above multilinear regressions for several functional sets of genes that were (a) enriched in the three clusters containing genes expressed at medium or high levels (clusters 6, 7 and 8, Figure 3), (b) involved in epigenetic regulation or neural function, and (c) relevant to male fly behavior. Since GO analysis is ineffective in functionally classifying small sets of genes, we manually categorized these genes in each group using their functions defined in Flybase. Genes in the following 9 functional groups were identified: sleep, neuropeptide, male mating, G-protein signaling, ligand-gated ion channel, catecholamine metabolism, MAPK signaling, and certain epigenetic genes (Table 2). The writers and erasers of marks were grouped by whether their marks tend to activate or repress gene transcription.

**Table 2:** mRNA changes between GH and SH males are predicted by changes in epigenetic marks and presence of some TF binding sites. The ability to predict is given by the coefficient of multiple correlation r, and the p-value from an F-test. Full statistics are given in Supplement Data 3. The sign of the partial correlation coefficient is given as +, -, or blank for non-significant values. (-) indicates the coefficient was marginally significant.

|  | *Sleep-related genes* | *Neuropeptides and receptor genes* | *Male mating genes* | *G-protein signaling genes* | *Ligand-gated ion channel genes* | *Catecholamine metabolism genes* | *MAPK signaling genes* | *Epigenetic activation genes* | *Epigenetic repression genes* |
| --- | --- | --- | --- | --- | --- | --- | --- | --- | --- |
| Group | 1 | 2 | 3 | 4 | 5 | 6 | 7 | 8 | 9 |
| r | 0.31 | 0.25 | 0.53 | 0.34 | 0.39 | 0.72 | 0.30 | 0.79 | 0.86 |
| p | 0.01 | 0.05 | 0.001 | 0.001 | 0.03 | 0.03 | .005 | 0.007 | 0.006 |
| Hr38 |  | - | - | (-) | - |  | - | - |  |
| Cbt | - |  | - |  |  |  |  |  | + |
| CrebA |  |  |  | - |  |  |  |  |  |
| Sr | + |  |  |  |  | - |  |  |  |
| Pho |  |  |  |  |  |  | + | + | + |

Interestingly, significant amounts of mRNA change between GH and SH flies were explained by TF binding sites in these functional groups, as shown in Table 2. The table shows the r and p-values for the regressions, and which TF motifs were significantly different. *Hr38, cbt, CrebA*, and *sr* putative binding sites each show a significant connection to mRNA change in one or more of the 6 functional gene groups. Of note, in eight out of nine functional groups where ARG-TFs motifs were significantly different, the direction of the effect of *Hr38*, *cbt*, and *CrebA* binding sites was negative – that is, the more potential binding sites the TF had in a gene, the lower the difference in mRNA between GH and SH flies was. This is consistent with the putative role of ARGs as transcriptional repressors for some genes. By contrast, *pho*, known primarily as a transcriptional repressor (Brown et al., 1998), showed a consistent positive effect on fold change between GH and SH flies.

In summary, changes in the numbers of a few putative transcription factor binding sites were sufficient to predict mRNA changes due to housing in ten functionally relevant gene groups with r values ranging from 0.25 to 0.86 (Table 2). The influence of ARGs on differential transcription in GH and SH males in biologically relevant gene groups led us to hypothesize that ARGs might also affect phenotypes known to vary with housing conditions.

### Regulation of social isolation-induced behavior by ARGs

Social isolation has a robust influence on behavior; for example, SH flies show reduced daytime sleep when compared to GH flies (Brown et al., 2017; Ganguly-Fitzgerald et al., 2006). Having shown a potential involvement of ARG-TFs in regulating some genes differentially-expressed in dopaminergic neurons of GH and SH males, we knocked-down expression of these ARG-TFs in *TH-GAL4* neurons and assayed the males for their sleep patterns. Specifically, we quantified the differences in sleep between GH and SH males (or ΔSleep as described by Ganguly *et al.* 2006) in which these ARGs and epigenetic modifiers were downregulated in dopaminergic neurons using RNA interference. Knockdown of all four ARG-TFs (*CrebA*, *Hr38*, *cbt*, and *sr*), significantly reduced ∆Sleep (Figure 5A-C, Supplement Data 4). These data show that these ARG-TFs play significant roles in regulating social effects on sleep in dopaminergic neurons.

**Figure 5:**
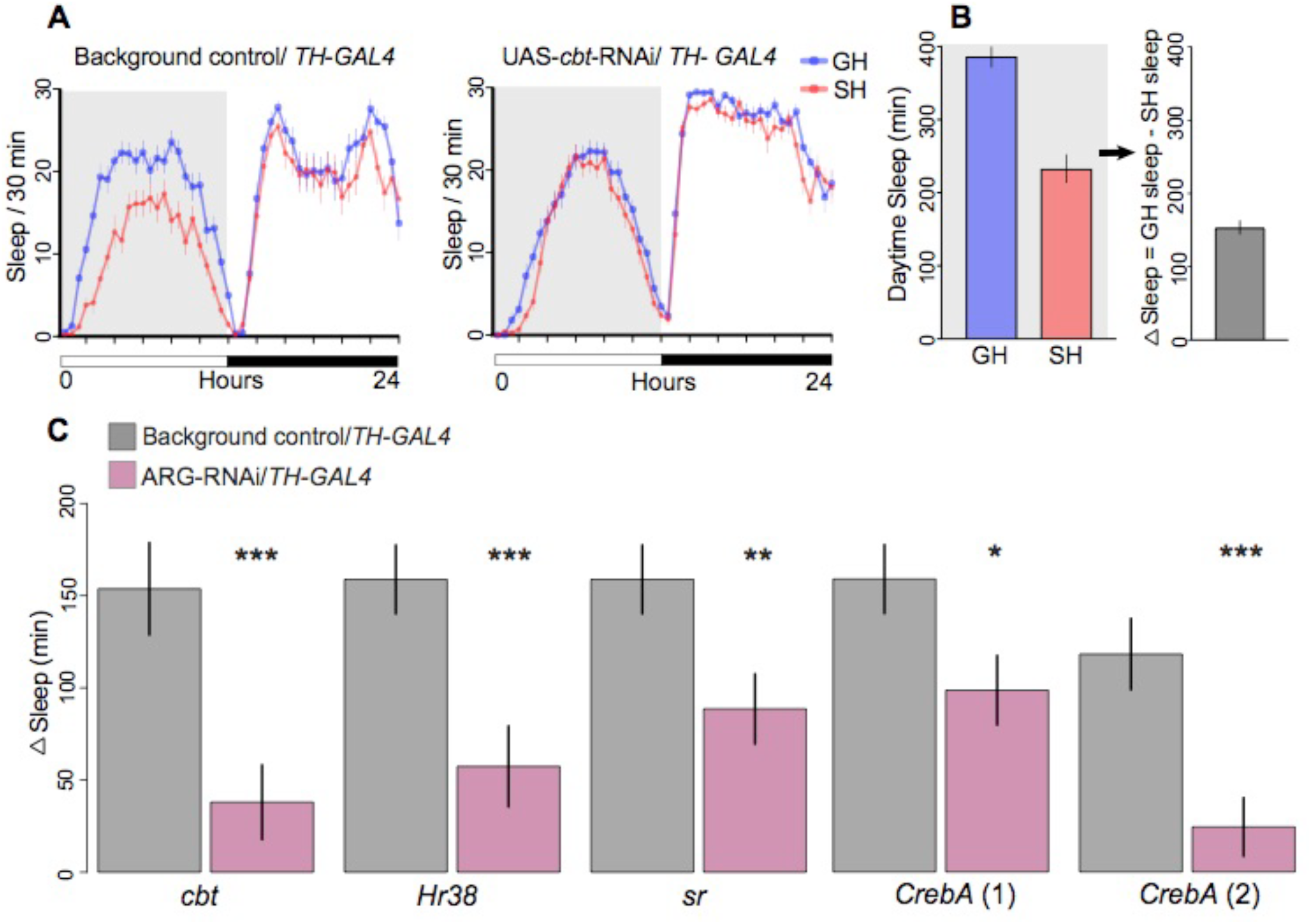
Knockdown of ARGs by RNAi affected social effects on daytime sleep. Knock-down of ARGs in dopaminergic neurons was achieved by driving RNAi transgenes with *TH-GAL4*; controls carried empty vectors without RNAi hairpin and *TH-GAL4*. (A) Example graph of sleep per 30 minutes over 24 hours. Control single housed (SH) flies sleep less than group housed (GH) flies during the day (shaded grey area). Expressing RNAi for ARG-TF *cabut* in dopaminergic neurons significantly reduced this difference. (B) Daytime sleep was measured and ΔSleep was compared between experimental males carrying the RNAi transgene and controls. ∆Sleep is defined as minutes of daytime sleep for GH flies minus the same measure for SH flies (as described by Ganguly *et al.* 2006). (C) ΔSleep for controls and RNAi knockdowns. Error bars are mean±SEM. In every case, RNAi knockdown significantly reduced the social effect on Δsleep. Two different RNAi lines were tested for CrebA, each showing significant reductions.

Our bioinformatic analysis suggested that these ARG-TFs might act as transcriptional repressors on downstream targets. Further analysis suggested that genes repressed by Cbt (Bartok et al., 2015) have reductions in H3K27ac and increases in H3K27me3 marks (Figure 4C). *Brms1* is a member of the histone deacetylase Sin3A repressor complex that contributes to PRC2 activity by deacetylating H3K27, thus allowing H3K27me3 to increase (Spain et al., 2010), and its transcription was also upregulated in GH flies. *Brms1* knockdown also significantly reduced ∆Sleep (Figure 5- Figure Supplement1), which is consistent with effects of Cbt on ∆Sleep (Figure 5C).

## Discussion

Results from this study provide insights into how social experiences, such as social isolation and social enrichment, can affect the epigenome of a small, well defined neural population in the adult *Drosophila* brain. We miniaturized the INTACT method, mini-INTACT, and examined epigenetic changes in a rare cell-type isolated from 200-250 adult fly heads. We carried out ChIP-seq on mini-INTACT purified dopaminergic neuronal nuclei for six different histone modification marks and correlated it to transcriptional profiles determined by RNA-seq. We found changes in the epigenetic landscape in dopaminergic neurons upon social experience in several gene-clusters. Our analysis identified four ARG-TFs (Chen et al., 2016) responding to social enrichment in dopaminergic neurons. RNAi-mediated knockdown of all four of these ARG-TFs (*cabut, Hr38, stripe, CrebA*) as well as an epigenetic eraser *Brms1* reduced the effects of social experience on daytime sleep (Figure 5, Supplement Data 4).

K-means clustering identified several differences in the epigenetic and transcriptional landscape that correlate with social experience. Curiously, many of the genes with highest mean mRNA expression levels also have higher levels of the repressive H3K27me3 mark and lower levels of some activating marks (Figure 3, cluster 8). In clusters of genes with lower expression levels, a more classical pattern of high levels of activating marks and low levels of repressive marks was found. But as the expression levels increase from these “classical” gene clusters towards the higher expression clusters (Figure 3, cluster 8), some activating marks drop and some repressive marks rise. In fully repressed genes, repressive PRC2-related H3K27me3 levels are uniformly higher than repressive PRC1-related H3K9me3 levels. However, in the transition from classical pattern of marks to the higher expression paradoxical pattern, H3K9me3 increases before H3K27me3.

Dopaminergic neurons in the fly brain are essential parts of circuits involved in learning and memory (Burke et al., 2012; Liu et al., 2012b; Waddell, 2013). Ganguly *et al.* showed that increased daytime sleep in GH males was associated with higher brain dopamine levels, and that it could be blocked by ablation of dopaminergic neurons or loss-of-function alleles of many learning and memory genes (Ganguly-Fitzgerald et al., 2006). Our finding that some genes highly expressed in dopaminergic neurons are associated with unusual pattern of epigenetic marks (Figure 3, cluster 7 and 8) is consistent with the finding that mouse differentiated dopaminergic neurons still contain substantial numbers of genes labelled "PRCa" or Polycomb Repressive Complex - Active, with both active transcription but also presence of repressive H3K27me3 marks (Ferrai et al., 2017). Another recent study found that in embryonic stem cells such PRCa genes have a higher variability of gene expression (Kar et al., 2017). Taken together, we suggest that some of the genes in fly dopaminergic neurons that show a change in expression between SH and GH males may be similar to those called PRCa genes in mouse dopaminergic neurons.

In the search for insect equivalents of immediate early genes, the Rosbash group identified ARGs in dopaminergic neurons (Chen et al., 2016). The fold change in response to stimulation in their top 50 ARGs correlates significantly with the fold changes we measured in mRNA in response to putative stimulation provided by group housing. The top 50 ARG fold changes in Chen *et al.* also correlate significantly with GH versus SH fold changes in H3K4me3, H3K9me3, and H3K27me3 in dopaminergic neurons in our study. Genes encoding four transcription factors (*CrebA, Hr38, sr* and *cbt)* are in the top genes by fold-change in both the Rosbash study (Chen et al., 2016), and in our own data.

*Hr38* and *stripe* have recently been shown to be activity-regulated in the honey bee and to affect dopamine pathway genes (Singh et al., 2018). *Hr38* is a homolog of vertebrate immediate early genes NR4A1-3, and has been shown to regulate dopaminergic neuron transcription and development (Eells et al., 2012; Kadkhodaei et al., 2009; Zetterström et al., 1996). In flies, *Hr38* overexpression increases dopamine decarboxylase (*Ddc*) transcription in the larval brain (Davis et al., 2007). In our data *Hr38* and *Ddc* are significantly higher in GH than in SH flies (Figure 2F and Figure 4A). Thus, although ARGs are co-expressed in a much broader range of neural types than just dopaminergic neurons (Croset et al., 2018) in the adult fly brain, they may have specific effects in dopaminergic neurons.

Cbt is a transcriptional repressor in some contexts, for instance in adult male fly heads (Bartok et al., 2015). Its vertebrate ortholog KLF10/TIEG1 acts with epigenetic repressors such as the H3K4 demethylase JARID1/KDM5B (Kim et al., 2010) and H3K27 deacetylase BRMS1 in the Sin3A complex (Belacortu et al., 2012; Muñoz-Descalzo et al., 2007; Spittau et al., 2007). Notably, if H3K27ac is deacetylated by BRMS1, this allows for the creation of the PRC2 mark H3K27me3 (Spain et al., 2010). We identified genes that were repressed or enhanced by Cbt and were in our top 40% expression range. These genes had highly significant differences in social-housing effects on mRNA and epigenetic mark levels, including downregulation in GH mRNA and H3K27ac and upregulation in GH H3K27me3 marks (Figure 4C).

RNAi knockdown of *Brms1* reduced the social housing effect on daytime sleep in the similar manner as *cbt* knockdown. Thus, we have a consistent picture in which genes repressed by Cbt (Bartok et al., 2015) have reductions in H3K27ac and increases in H3K27me3 marks (Figure 4C), and knockdown of the deacetylase for H3K27 produce effects on sleep similar to *cbt* (Figure 5- Figure Supplement 1). Further studies are needed to elucidate possible epigenetic pathways mediated by *CrebA, Hr38,* and *stripe.* Croset *et al.* suggest that the highly inter-correlated set of ARGs they found may have repressive effects on transcription in various brain regions, such as the mushroom body γ lobes (Croset et al., 2018). We found in our data that upregulation of some these genes (especially *cbt* and *Hr38*) in GH males is associated with downregulation of genes in some functional gene groups. These groups fall largely into the k-means cluster that has increases in PRC2-related marks (H3K27me3) in *TH-GAL4*-expressing dopaminergic neurons. This suggests a scenario where group housing stimulates ARG expression, and these TFs in turn down-regulate neural function genes in part by increasing PRC2 repressive marks.

*Drosophila* has been a successful model for neuro-genetics due to ease of manipulating flies, availability of large collection of genetic tools, and recent development of automated behavioral assays. Adaptation of cell-type specific epigenetic methods such as mini-INTACT can help leverage this potential to comprehensively study epigenetic changes in specific neurons across several paradigms including stress, drugs of abuse, neuro-degenerative disorders etc.

## Materials and Methods

### Fly stocks & rearing

*Drosophila melanogaster* in a Canton-S background were reared on standard fly food at 25^°^C at 65% relative humidity with a 12/12 h light/dark cycle. For social isolation and group housing experiments 24-48 hour-old males of a given genotype were housed individually or in groups of 20 in standard *Drosophila* vials (2.6 cm diameter X 9.3 cm high) for 4 days containing standard fly food. *3X*-, *5X*- and *10X-UAS-unc84-2XGFP* and *10XUAS_unc84-tdTomfl* are as described (Henry et al., 2012) and were a kind gift of Henry Gilbert (Janelia Research Campus, VA, USA), *TH-GAL4* is as described (Friggi-Grelin et al., 2003). Tissue collections for genomic analysis were performed near morning activity peak, usually around ZT3-ZT5. The following TRiP RNAi lines (Perkins et al., 2015) were obtained from the Bloomington Stock Center for behavioral analysis: BL36303 (*y[1] v[1]; P{y[+t7.7]=CaryP}attP2*) no insert background control *vs.* RNAi lines: BL29377 (*Hr38*); BL31900 (*CrebA*); BL27701 (*Sr*). BL36304 (*y[1] v[1]; P{y[+t7.7]=CaryP}attP40*) no insert background control *vs.* RNAi lines: BL42562 (*CrebA*); BL38276 (*cbt*) and BL42533 (*Brms1*).

### Immunostaining and imaging

Fly brains were dissected in cold 1X phosphate buffered saline (PBS) and fixed in 2% paraformaldehyde made in 1X PBS at room temperature for 1 h on a nutator, washed 4 times for 20 min each in PAT (1X PBS, 0.5% PBS Triton, 1% BSA) at room temperature, blocked for 1hour at room temperature with blocking buffer (PAT + 3% Normal Goat Serum) and incubated with primary antibodies, diluted in blocking buffer, overnight on a nutator at 4°C. The primary antibodies used were: Mouse-GFP (SIGMA-ALDRICH, G6539. 1:500 dilution), Rabbit-TH (EMD-Millipore, AB152, 1:200 dilution), and Rat-DN-cadherin (Hybridoma Bank DSHB, DNEX#8, 1:50 dilution). This was followed by 4 washes for 20 min each in PAT, and incubation overnight on a nutator at 4°C with secondary antibodies diluted in blocking buffer. The secondary antibodies were all from Molecular Probe and used at 1:500 dilution: Alexa Fluor 488 anti-Mouse (A11029), Alexa Fluor 568 anti-Rabbit (A11036) and Alexa Fluor 633 anti-Rat (A21094). Brains were then washed 4 times for 20 min each in PAT at room temperature, 1 time for 20 min in 1X PBS and mounted with VECTASHIELD mounting medium (Vector Laboratories, H-1000). Samples were imaged on a Zeiss 800 confocal laser-scanning microscope.

### mini-INTACT

Nuclei were obtained from dopaminergic neurons using INTACT (Henry et al., 2012) with modifications to enable purification of nuclei from as few as 200-250 heads per ChIP-seq for *TH*-*GAL4* which is expressed in ~120 neurons/brain. *Drosophila* males of *3X-UAS-unc84-2XGFP/TH-GAL4* genotype were either socially isolated or group housed and flash frozen during the morning activity peak. Frozen heads were collected over dry ice-cooled sieves from vortex-decapitated flies and added to 5 ml of mini-INTACT buffer consisting of 5mM β-glycerophosphate pH 7.0, 2 mM MgCl2, 1x complete protease inhibitor cocktail (Roche: 11873580001), 5 mM sodium butyrate, 0.6 mM spermidine, 0.2 mM spermine, 0.5% NP-40 and 0.6mM β-mercaptoethanol. The suspension was passed over a continuous flow homogenizer, set at 1000 rpm, ten to twelve times. The homogenizer was modified such that the grooves at the bottom of the homogenizer helped push fly heads upward increasing efficiency of homogenization and preventing sample loss (Figure 1- Figure Supplement 1). Homogenate was filtered through a 20 μm filter (Partec CellTrics, Sysmex: 25004-0042-2315) and then a 10 μm filter (Partec CellTrics, Sysmex: 04-0042-2314). 1 μg of anti-GFP antibody (Invitrogen: G10362) was added to the filtered homogenate, tubes were gently inverted 10 times, and incubated on ice for 20 minutes to allow binding. To this mix, 30 μl of Dynabeads Protein-G (Invitrogen: 100-03D) were added and incubated at 4°C for 30 min with constant end-over-end rotation. Beads were then collected on a magnet (Diagenode: B04000003) and washed thrice using INTACT buffer. Bead-bound nuclei were resuspended in 1 ml INTACT buffer and formaldehyde fixed for ChIP-seq as described in the next section.

### ChIP-Seq

For each ChIP-seq reaction ~10,000-15,000 mini-INTACT isolated bead-bound nuclei were processed using Low Cell # ChIP kit (Diagenode: C01010070) as per manufacturer’s instructions. In brief, nuclei were fixed in 1% formaldehyde for 2 minutes, immediately quenched with Glycine and then lysed using nuclear lysis buffer with protease inhibitor cocktail at room temperature for 5 minutes. PBS was added to dilute the lysate-bead mix and loaded in AFA tubes (Covaris Inc.: 520045) for sonication. Ultra-sonicator (Covaris Inc.: E220) was used to sheer chromatin to ~200 bp length and chromatin was recovered from the supernatant after magnetic separation. ChIP was performed using the following ChIP-seq grade antibodies: H3K4me3 (Diagenode: C15410003-50), H3K9me3 (Diagenode: C15410193), H3K9/K14ac (Diagenode: C15410200), H3K27me3 (Diagenode: C15410195), H3K27ac (Diagenode: C154410196) and H3K36me3 (Diagenode: C15410192). Two biological replicates were performed for each histone mark and input DNA was used as the control. Libraries for sequencing were prepared using MicroPlex Library Preparation kit (Diagenode: C05010012) as per manufacturer’s instruction. Single end 60 bp sequencing reads were obtained using Illumina Hi-seq 2500.

### RNA-seq

We isolated cell bodies of dopaminergic neurons using Fluorescence Activated Cell Sorting (FACS) during the flies’ morning activity peak. The protocol was essentially as described (Hempel et al., 2007) with minor modifications. In brief, brains were dissected from socially-isolated or group-housed flies expressing membrane-tagged GFP and nuclear tdTomato in their dopaminergic neurons. The flies were obtained by crossing flies carrying *TH-GAL4* with a stock carrying *pJFRC105-10XUAS-IVS-nlstdTomato* in VK40 (gift of Barret D. Pfeiffer, Rubin Lab, Janelia Research Campus) and *pJFRC29-10XUAS-IVS-myr::GFP-p10* in AttP40 (Pfeiffer et al., 2012) and was found to produce better purity in FACS than other reporters (Etheredge et al., 2018). To account for possible manual bias, dissectors switched their handling of group- or single-housed flies in each session. Dissected brains were digested using Liberase DH (Roche: 5401054001), manually triturated using glass pipettes, and filtered using a Falcon 35 μm cell strainer (Corning: 352235) before sorting. Approximately 1500 dopaminergic neurons were obtained from approximately 30 brains using a BD FACSAria II sorter (BD Biosciences, USA). Total RNA was extracted using the Arctus, PicoPure RNA Isolation Kit (Thermo Fisher Scientific: 12204-01), ERCC spike-in controls were added and cDNA libraries from this material were prepared using Ovation RNA-seq System V2 (Nugen: 7102) as per manufacturer’s instructions. Three biological replicates were performed for each condition. Paired end 100 bp sequencing reads were obtained using Illumina Hi-seq 2500.

### Sleep assay

Flies that were previously socially isolated or group housed for 4 days were anesthetized briefly with carbon dioxide and transferred into 5 mm × 65 mm transparent plastic tubes with standard cornmeal dextrose agar media. For recording locomotor activity, *Drosophila* activity monitors (Trikinetics, Waltham, USA) were kept in incubators at 25^°^C with 65% relative humidity in a 12/12 h light/dark cycle. Flies were allowed one night to acclimatize to the apparatus and activity data was collected in 1 minute bins for the following 24 hours as described (Donelson et al., 2012). One sleep bout was defined as 5 minutes of continuous inactivity (Hendricks et al., 2000; Shaw et al., 2000). Statistical analysis of the sleep data was performed using Prism 7 (GraphPad software) and R scripts (R Core Team, 2014).

### Bioinformatics

#### Sequencing analysis

All genomic procedures used release 6.02 of the *Drosophila melanogaster* genome (Dos Santos et al., 2015). R 3.0.3 was used in scripts and statistics (R Core Team, 2014). Non-parametric statistical tests were used except where noted. STAR (Dobin et al., 2013) was used for alignment of RNA-seq data. Total counts of de-duplicated reads were calculated at each genome position using Rsubread (Liao et al., 2013), followed by differential expression calls using edgeR (Robinson et al., 2010). We cross-checked differential expression using the CyberT (Kayala and Baldi, 2012) and FCROS (Dembélé and Kastner, 2014) packages. Normalization between replicates and treatments was performed using default methods (TMM) in edgeR to correct for coverage levels. CuffDiff (Trapnell et al., 2012) was used to detect changes in splicing. Bowtie (Langmead, 2010) was used to align ChIP-seq reads, and DiffReps (Shen et al., 2013) and ngs.plot (Shen et al., 2014) were used to quantify ChIP-seq reads. Changes to DiffReps and ngs.plot databases and code were required to use *Drosophila* genome release 6.02 and are included in Supplement Data 5. SICER (Xu et al., 2014a; Zang et al., 2009) was also run to cross-check DiffReps results (Supplement Table 6).

### Clustering

Gene Ontology (GO) analysis was done using two web tools: DAVID (Huang et al., 2009) and GOrilla (Eden et al., 2009). For mRNA differential expression analysis, genes in the top 40% of expression level were used as the background lists for both tools, and genes with FCROS significant differential expression (FDR=0.2) were analyzed. For analysis of clusters (see below) genes belonging to each cluster were compared to the appropriate background list (top 40% genes for 8-clusters). K-means clustering (Hartigan and Wong, 1979) was done using the kmeans package in R. To understand the impact of social isolation on epigenetics of genes expressed in dopaminergic neurons, a dataset of the top 40% of genes by mRNA TPM expression (5,372 genes) was constructed containing normalized differences between group-housed and isolated flies for mRNA and for the 6 epigenetic marks. Tests using an information criterion approach (BIC) were used to determine the optimal numbers of clusters, which was k=8 for the 5,372-gene dataset. K-means clustering is a stochastic process that may yield very different results each time it is run if there is no strong pattern in the data. To determine robustness of the gene assignments to clusters, we re-ran clustering with random seed changes to create N cluster assignments. We then compared each cluster assignment to every other (1035 = N*(N-1)/2 comparisons for 8-cluster assignments). In each comparison, we calculated the percent overlap of a cluster in assignment i with clusters in assignment j, and reported the maximum percent overlap for that cluster. We therefore generated 8,280=8*1,035 comparisons. Figure 3- Figure Supplement 1 shows a histogram of cluster overlap percentages. For eight clusters, the median percent overlap of a cluster in one assignment to its best match in a second assignment was 94%, and was >99%, 72% of the time. Thus, we concluded that cluster identity is fairly stable, in spite of the randomness inherent in the k-means algorithm.

Cluster functional enrichment was determined using the DAVID 6.8 functional annotation tool (Huang et al., 2009) using biological, cellular, and molecular function levels 5 plus chromosome location, and using functional annotation clustering. For the gene clusters the GO analysis by DAVID used the 5,372 highest expression genes as background. Results are reported using thresholds for individual categories FDR < 0.05 and enrichment value >2.0 for functional clusters.

### Motif analysis

We used the MEME suite of tools (Bailey et al., 2009) to find putative transcription factor binding sites in promoters of the 8 gene clusters found by k-means. Centrimo 4.12.0 (Bailey and Machanick, 2012) was used with promoter-proximal (± 500 bp from TSS) sequences of genes. We used databases of TF binding motifs from Fly Factor Survey 2014 (Enuameh et al., 2013; Zhu et al., 2011) supplemented by motifs determined in a recent study (Nitta et al., 2015). Promoter proximal sequences of each gene in a cluster (“test genes”) were tested for motif enrichment using Centrimo compared to a control set of sequences from an equal number of randomly selected genes not in the cluster (“control genes”). We report a motif as “enriched” if the Centrimo’s adjusted p-value was < 1 × 10^−10^.

To quantify the number of potential binding sites of each enriched motif in each gene, we used FIMO version 4 (Grant et al., 2011) with default parameters. Log fold changes in mRNA levels between group housed and single housed treatments were the dependent variable in multilinear regressions in which numbers of each enriched TF motifs were used as dependent variables. The “lm” program from R was used; non-significant dependent variables were removed in a step-wise manner using “stepAIC” (least significant first) until only significant variables remained; the results of these regressions are reported with multilinear r (square root of proportion of variance explained by the regression) and F-test p-value. Full tables of regression fits are provided in Supplement Data 3.

## Acknowledgements

We thank Lee Henry Gilbert for help with INTACT protocol and fly stocks, Janelia Cell Culture facility for help with FACS, Igor Negrashow and Janelia Experimental Technology for help designing the homogenizer, Serge Picard and Janelia Quantitative Genomics for sequencing and Andrew Lemire for helpful discussions. We thank Barret Pfeiffer, Gerald Rubin and Jack Etheredge for fly stocks used in FACS. We would also like to thank Loren Looger and members of Heberlein lab for helpful discussions and Mark Cembrowski and Vilas Menon for comments on the manuscript. This work was supported by the Howard Hughes Medical Institute.

## Tables and Figures with legends

**Figure 1- Figure Supplement 1:**
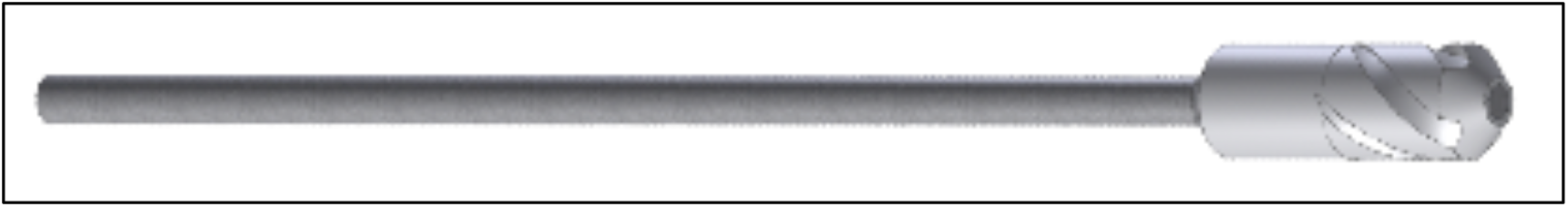
Design of homogenizer used in mini-INTACT.

**Figure 1- Figure Supplement 2:**
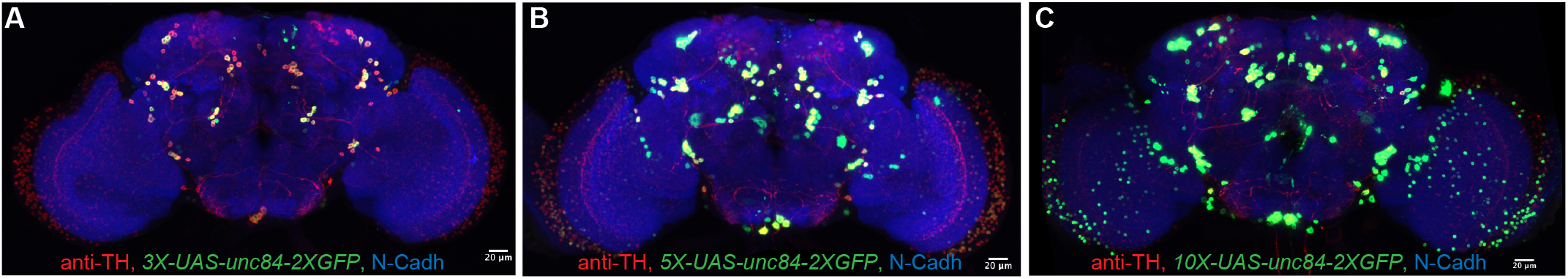
Comparison of tagged GFP expression in adult *Drosophila* brain. The INTACT transgene (*unc84-2XGFP*) was driven in dopaminergic neurons (TH*-GAL4*) using different copy numbers of the UAS promoter and expression of GFP was compared using the same imaging settings. (A) *3X-UAS*- (B) *5X-UAS*- and (C) *10X-UAS-unc84-2XGFP*. The *3X-UAS-unc84-2XGFP* transgene most faithfully reproduced *TH*-*GAL4* expression, while ectopic expression was observed upon further increases of the *UAS* copy numbers. Dopaminergic neurons were stained with anti-TH antibodies (red), INTACT transgene expression using anti-GFP antibodies (green), and N-cadherin (blue) was used as reference. See Figure 1B for *3X-UAS-unc84-2XGFP* brain imaged at higher intensity. Scale bar is 20 μm.

**Figure 1- Figure Supplement 3:**
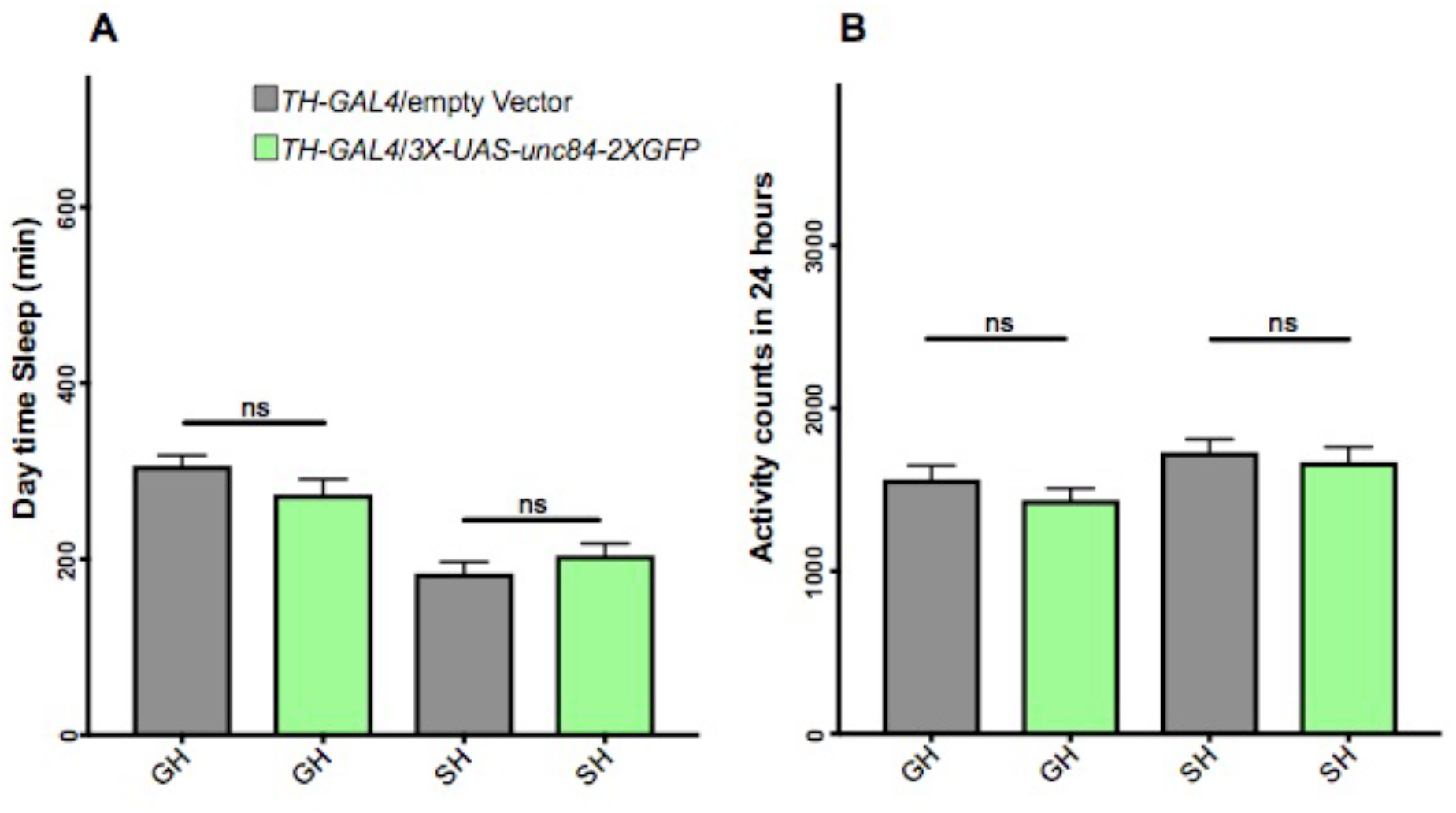
*3XUAS-unc84-2XGFP* expression in dopaminergic neurons did not affect daytime sleep or activity over 24 hours. (A) Daytime sleep measured over a 12 hours period. GH males sleep more than SH males during the daytime. No significant difference was observed due to tagged-GFP expression. (B) Total number of activity counts (beam breaks) over 24 hours. GH are less active than SH flies as expected. No significant difference was observed due to tagged-GFP expression. N = 31-32. Unpaired t-test.

**Figure 1- Figure Supplement 4:**
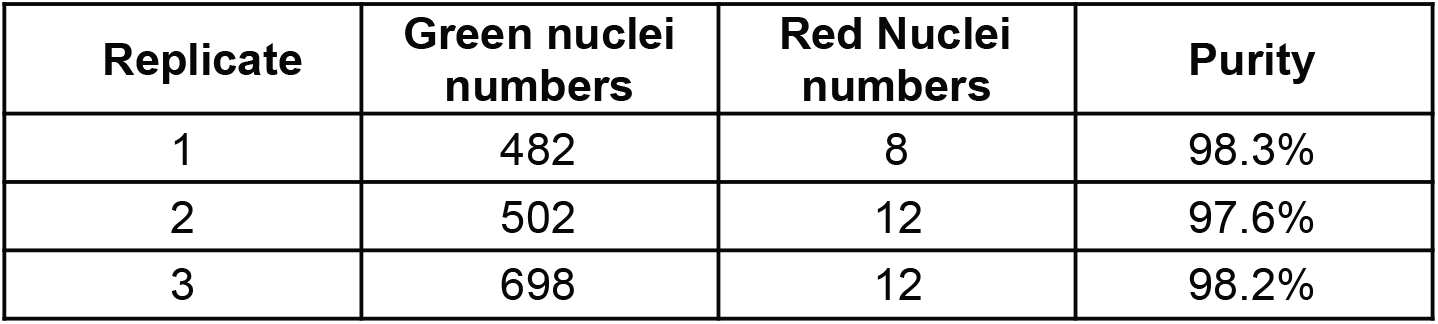
Purity assessment of dopaminergic nuclei. The table shows the number of captured green dopaminergic nuclei using bead-bound anti-GFP antibodies. Most of the contaminating red nuclei were washed away from bead-bound affinity-purified nuclei. See Figure 1 and main text for details (3 biological replicates).

**Figure 2- Figure Supplement 1:**
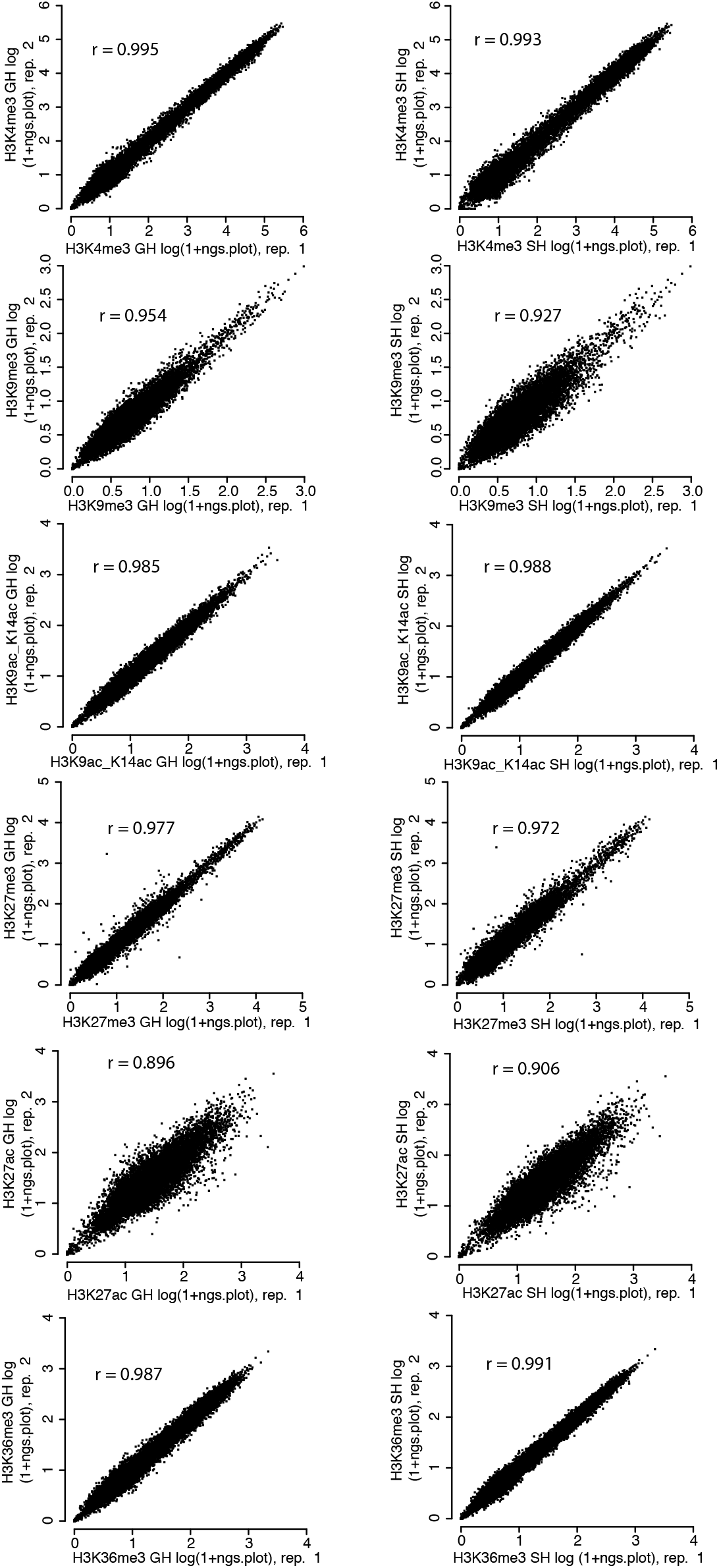
Replicate concordance for ChIP-seq for various histone modifications. ChIP-671 seq replicate concordance is shown with Pearson’s correlation coefficient (r-values) calculated on Log (1+ngs.plot) 672 enrichment values for all six histone marks.

**Figure 3- Figure Supplement 1:**
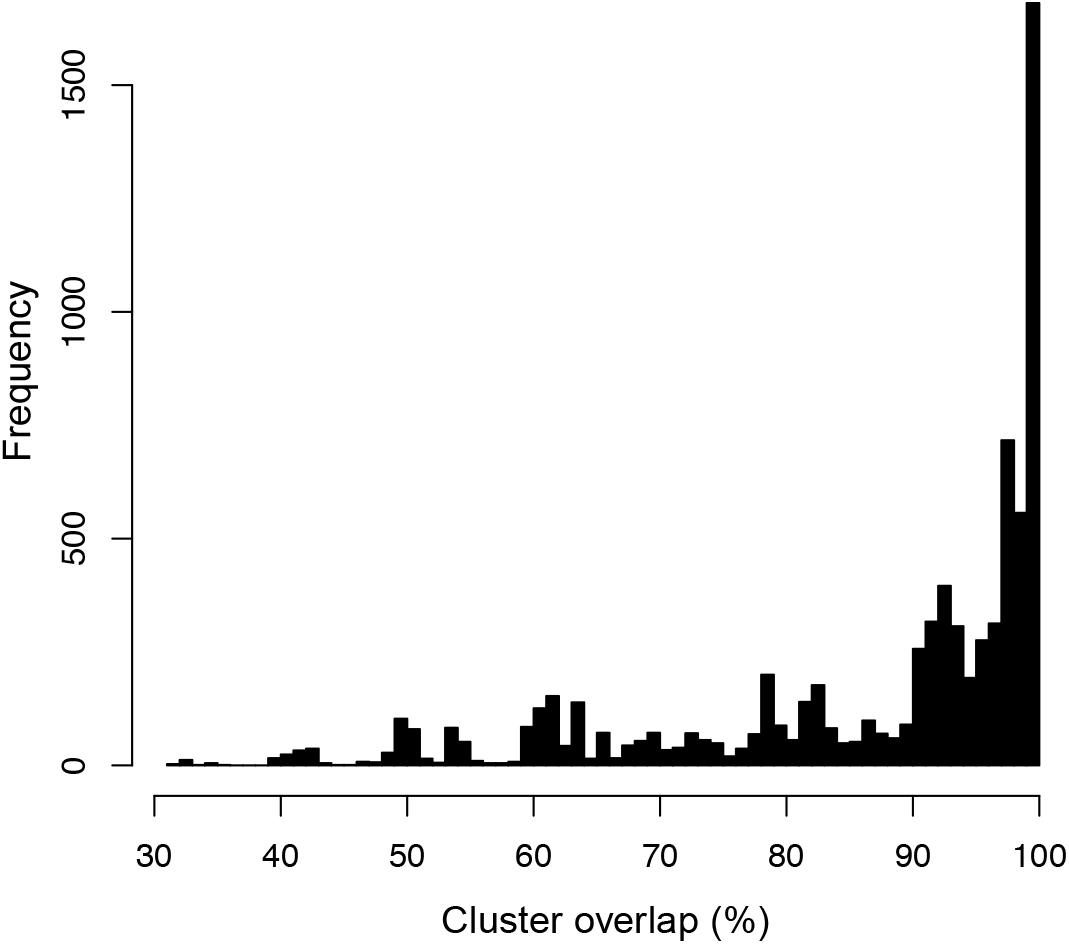
k-means cluster overlap. The figure shows a histogram of k-means cluster overlap percentages used to calculate robustness of gene assignments to clusters. For eight clusters, the median percent overlap of a cluster in one assignment to its best match in a second assignment was 94%, and was greater than 99% 72% of the time (see Methods for details).

**Figure 5- Figure Supplement 1:**
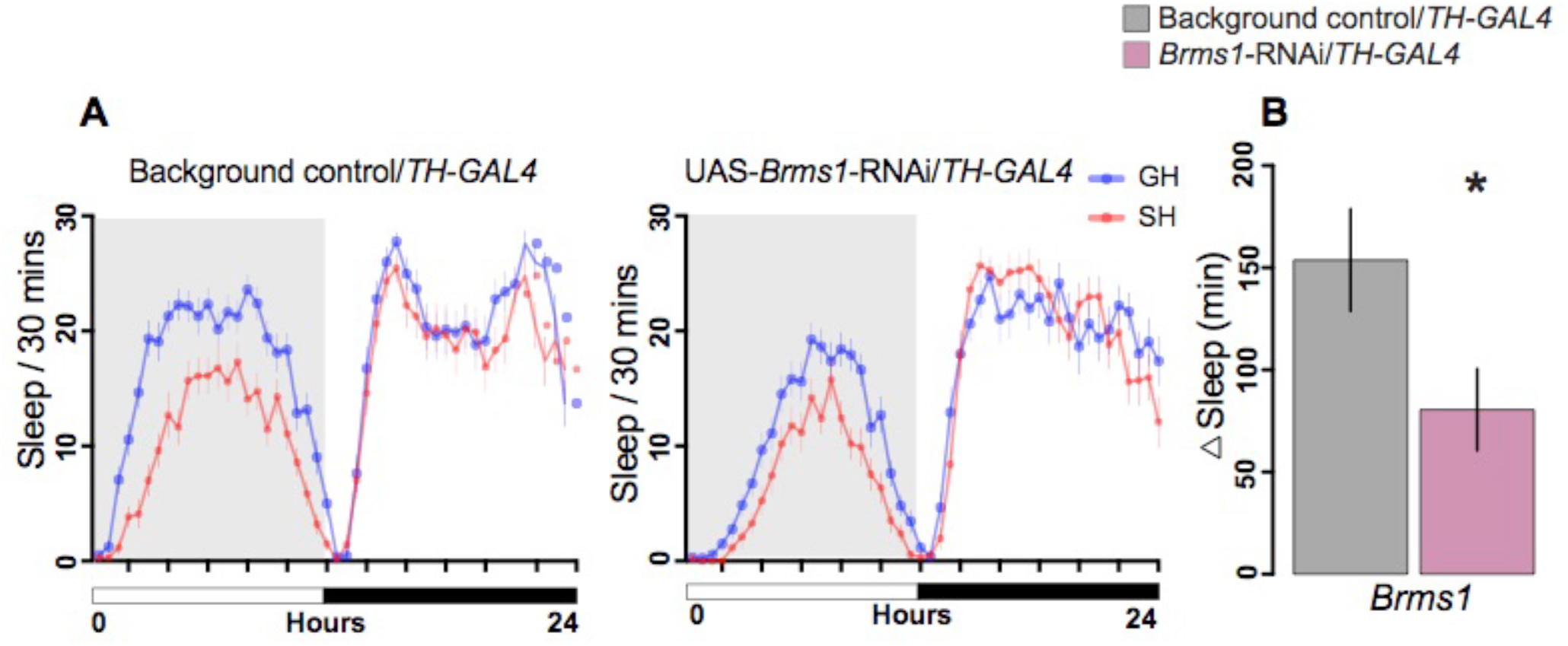
Knockdown of epigenetic eraser *Brms1* by RNAi reduced social effects on daytime sleep. *Brms1* is a member of *Sin3A* histone deacetylase complex. Knockdown of *Brms1* in dopaminergic neurons was achieved by driving an RNAi transgene with *TH-GAL4*; controls carried empty vectors without RNAi hairpin and *TH-GAL4*. (A) Sleep per 30 minutes over 24 hours for control and *Brms1* knockdown in SH and GH flies. Daytime sleep is highlighted in shaded grey area for both genotypes. (B) Expressing RNAi for *Brms1* in dopaminergic neurons reduced the social effect of sleep during the day (∆Sleep). Error bars are mean ±SEM.

## References

Abdelmoez, M. N., Iida, K., Oguchi, Y., Nishikii, H., Yokokawa, R., Kotera, H., Uemura, S., Santiago, J. G. and Shintaku, H. (2018). SINC-Seq:Correlation of gene expressions between nucleus and cytoplasm reflects single-cell physiology. Genome Biol., 19, 1–11.

Abruzzi, K. C., Zadina, A., Luo, W., Wiyanto, E., Rahman, R., Guo, F., Shafer, O. and Rosbash, M. (2017). RNA-seq analysis of Drosophila clock and non-clock neurons reveals neuron-specific cycling and novel candidate neuropeptides. PLoS Genet., 13, 1–23.

Alekseyenko, O. V, Chan, Y.-B., Li, R. and Kravitz, E. A. (2013). Single dopaminergic neurons that modulate aggression in Drosophila. Proc. Natl. Acad. Sci. U. S. A., 110, 6151–6.

Amin, N. M., Greco, T. M., Kuchenbrod, L. M., Rigney, M. M., Chung, M.-I., Wallingford, J. B., Cristea, I. M. and Conlon, F. L. (2014). Proteomic profiling of cardiac tissue by isolation of nuclei tagged in specific cell types (INTACT). Development, 141, 962–973.

Anreiter, I., Kramer, J. M. and Sokolowski, M. B. (2017). Epigenetic mechanisms modulate differences in Drosophila foraging behavior. Proc. Natl. Acad. Sci. U.S.A., 114, 12518–12523.

Anreiter, I., Biergans, S. D. and Sokolowski, M. B. (2019). Epigenetic regulation of behavior in Drosophila melanogaster. Curr. Opin. Behav. Sci., 25, 44–50.

Azanchi, R., Kaun, K. R. and Heberlein, U. (2013). Competing dopamine neurons drive oviposition choice for ethanol in Drosophila. Proc. Natl. Acad. Sci., 110, 21153–21158.

Bailey, T. L. and Machanick, P. (2012). Inferring direct DNA binding from ChIP-seq. Nucleic Acids Res., 40, e128.

Bailey, T. L., Boden, M., Buske, F. A., Frith, M., Grant, C. E., Clementi, L., Ren, J., Li, W. W. and Noble, W. S. (2009). MEME Suite: Tools for motif discovery and searching. Nucleic Acids Res., 37, 202–208.

Bainton, R. J., Tsai, L. T., Singh, C. M., Moore, M. S., Neckameyer, W. S. and Heberlein, U. (2000). Dopamine modulates acute responses to cocaine, nicotine and ethanol in Drosophila. Curr. Biol., 10, 187–94.

Barski, A., Cuddapah, S., Cui, K., Roh, T. Y., Schones, D. E., Wang, Z., Wei, G., Chepelev, I. and Zhao, K. (2007). High-Resolution Profiling of Histone Methylations in the Human Genome. Cell, 129, 823–837.

Bartok, O., Teesalu, M., Ashwall-Fluss, R., Pandey, V., Hanan, M., Rovenko, B. M., Poukkula, M., Havula, E., Moussaieff, A., Vodala, S., et al. (2015). The transcription factor Cabut coordinates energy metabolism and the circadian clock in response to sugar sensing. EMBO J., 34, 1538–1553.

Belacortu, Y., Weiss, R., Kadener, S. and Paricio, N. (2012). Transcriptional activity and nuclear localization of cabut, the Drosophila ortholog of vertebrate TGF-β-Inducible Early-Response gene (TIEG) proteins. PLoS One, 7, e32004.

Brand, A. H. and Perrimon, N. (1993). Targeted gene expression as a means of altering cell fates and generating dominant phenotypes. Development, 118, 401–15.

Brown, J. L., Mucci, D., Whiteley, M., Dirksen, M. L. and Kassis, J. A. (1998). The Drosophila polycomb group gene pleiohomeotic encodes a DNA binding protein with homology to the transcription factor YY1. Mol. Cell, 1, 1057–1064.

Brown, M. K., Strus, E. and Naidoo, N. (2017). Reduced sleep during social isolation leads to cellular stress and induction of the unfolded protein response. Sleep, 40, 7.

Burke, C. J., Huetteroth, W., Owald, D., Perisse, E., Krashes, M. J., Das, G., Gohl, D., Silies, M., Certel, S. and Waddell, S. (2012). Layered reward signalling through octopamine and dopamine in Drosophila. Nature, 492, 433–437.

Cacioppo, J. T., Ernst, J. M., Burleson, M. H., McClintock, M. K., Malarkey, W. B., Hawkley, L. C., Kowalewski, R. B., Paulsen, A., Hobson, J. A., Hugdahl, K., et al. (2000). Lonely traits and concomitant physiological processes: The MacArthur social neuroscience studies. Int. J. Psychophysiol., 35, 143–154.

Champagne, F. A. (2010). Epigenetic influence of social experiences across the lifespan. Dev. Psychobiol., 52, 299–311.

Chase, K. A. and Sharma, R. P. (2013). Nicotine induces chromatin remodelling through decreases in the methyltransferases GLP, G9a, Setdb1 and levels of H3K9me2. Int. J. Neuropsychopharmacol., 16, 1129–38.

Chen, X., Rahman, R., Guo, F. and Rosbash, M. (2016). Genome-wide identification of neuronal activity-regulated genes in Drosophila. Elife, 5, e19942.

Cirelli, C. and Tononi, G. (1998). Differences in gene expression between sleep and waking as revealed by mRNA differential display. Mol. Brain Res., 56, 293–305.

Croset, V., Treiber, C. D. and Waddell, S. (2018). Cellular diversity in the Drosophila midbrain revealed by single-cell transcriptomics. Elife, 7, e34550.

David, A. C., Marie-Laure, C., Thomas, J. and Reinberg, D. (2015). Epigenetics, 2nd Edition. Cold Spring Harbor Laboratory Press.

Davis, M. M., Yang, P., Chen, L., O’Keefe, S. L. and Hodgetts, R. B. (2007). The orphan nuclear receptor DHR38 influences transcription of the DOPA decarboxylase gene in epidermal and neural tissues of Drosophila melanogaster. Genome, 50, 1049–1060.

Deal, R. B. and Henikoff, S. (2010). A simple method for gene expression and chromatin profiling of individual cell types within a tissue. Dev. Cell, 18, 1030–1040.

Dembélé, D. and Kastner, P. (2014). Fold change rank ordering statistics: a new method for detecting differentially expressed genes. BMC Bioinformatics, 15,.

Dobin, A., Davis, C. a., Schlesinger, F., Drenkow, J., Zaleski, C., Jha, S., Batut, P., Chaisson, M. and Gingeras, T. R. (2013). STAR: ultrafast universal RNA-seq aligner. Bioinformatics, 29, 15–21.

Donelson, N., Kim, E. Z., Slawson, J. B., Vecsey, C. G., Huber, R. and Griffith, L. C. (2012). High-resolution positional tracking for long-term analysis of Drosophila sleep and locomotion using the “tracker” program. PLoS One, 7, e37250.

Dos Santos, G., Schroeder, A. J., Goodman, J. L., Strelets, V. B., Crosby, M. A., Thurmond, J., Emmert, D. B., Gelbart, W. M., Brown, N. H., Kaufman, T., et al. (2015). FlyBase: Introduction of the Drosophila melanogaster Release 6 reference genome assembly and large-scale migration of genome annotations. Nucleic Acids Res., 43, D690–D697.

Eden, E., Navon, R., Steinfeld, I., Lipson, D. and Yakhini, Z. (2009). GOrilla: A tool for discovery and visualization of enriched GO terms in ranked gene lists. BMC Bioinformatics, 10, 48.

Eells, J. B., Wilcots, J., Sisk, S. and Guo-Ross, S. X. (2012). NR4A gene expression is dynamically regulated in the ventral tegmental area dopamine neurons and is related to expression of dopamine neurotransmission genes. J. Mol. Neurosci., 46, 545–553.

Emmert-Buck, M. R., Bonner, R. F., Smith, P. D., Chuaqui, R. F., Zhuang, Z., Goldstein, S. R., Weiss, R. A. and Liotta, L. A. (1996). Laser Capture Microdissection. Science, 274, 998–1001.

Enuameh, M. S., Asriyan, Y., Richards, A., Christensen, R. G., Hall, V. L., Kazemian, M., Zhu, C., Pham, H., Cheng, Q., Blatti, C., et al. (2013). Global analysis of Drosophila Cys2-His2 zinc finger proteins reveals a multitude of novel recognition motifs and binding determinants. Genome Res., 23, 928–940.

Etheredge, J., Baumann, A. and Truman, J. W. (2018). Fluorescent reporter combination optimization for flow cytometry purity of labeled Drosophila neurons. https://doi.org/10.6084/m9.figshare.6934250.v1.

Febinger, H. Y., George, A., Priestley, J., Toth, L. A. and Opp, M. R. (2014). Effects of housing condition and cage change on characteristics of sleep in mice. J. Am. Assoc. Lab. Anim. Sci., 53, 29–37.

Feng, J., Wilkinson, M., Liu, X., Purushothaman, I., Ferguson, D., Vialou, V., Maze, I., Shao, N., Kennedy, P., Koo, J., et al. (2014). Chronic cocaine-regulated epigenomic changes in mouse nucleus accumbens. Genome Biol., 15, R65.

Ferguson, C. J., Averill, P. M., Rhoades, H., Rocha, D., Gruber, N. P. and Gummattira, P. (2005). Social isolation, impulsivity and depression as predictors of aggression in a psychiatric inpatient population. Psychiatr. Q., 76, 123–137.

Ferrai, C., Torlai Triglia, E., Risner-Janiczek, J. R., Rito, T., Rackham, O. J., de Santiago, I., Kukalev, A., Nicodemi, M., Akalin, A., Li, M., et al. (2017). RNA polymerase II primes Polycomb-repressed developmental genes throughout terminal neuronal differentiation. Mol. Syst. Biol., 13, 946.

Fitzsimons, H. L., Schwartz, S., Given, F. M. and Scott, M. J. (2013). The histone deacetylase HDAC4 regulates long-term memory in Drosophila. PLoS One, 8, e83903.

Friedman, E. M. (2011). Sleep quality, social well-being, gender, and inflammation: An integrative analysis in a national sample. Ann. N. Y. Acad. Sci., 1231, 23–34.

Friggi-Grelin, F., Coulom, H., Meller, M., Gomez, D., Hirsh, J. and Birman, S. (2003). Targeted gene expression in Drosophila dopaminergic cells using regulatory sequences from tyrosine hydroxylase. J. Neurobiol., 54, 618–27.

Ganguly-Fitzgerald, I., Donlea, J. and Shaw, P. J. (2006). Waking experience affects sleep need in Drosophila. Science, 313, 1775–81.

Ghezzi, A., Krishnan, H. R., Lew, L., Prado, F. J., Ong, D. S. and Atkinson, N. S. (2013). Alcohol-induced histone acetylation reveals a gene network involved in alcohol tolerance. PLoS Genet., 9, e1003986.

Gozen, O., Balkan, B., Yildirim, E., Koylu, E. O. and Pogun, S. (2013). The epigenetic effect of nicotine on dopamine D1 receptor expression in rat prefrontal cortex. Synapse, 67, 545–52.

Grant, C. E., Bailey, T. L. and Noble, W. S. (2011). FIMO: Scanning for occurrences of a given motif. Bioinformatics, 27, 1017–1018.

Greco, A. M., Gambardella, P., Sticchi, R., Federico, N. and Pansini, V. S. (1988). Chronic Administration of Imipramine Antagonizes Deranged Circadian Rhythm Phases in Individually Housed Rats. Physiol. Behav., 67–72.

Grippo, A. J., Cushing, B. S. and Carter, C. S. (2007). Depression-like behavior and stressor-induced neuroendocrine activation in female prairie voles exposed to chronic social isolation. Psychosom Med, 69, 149–157.

Gupta, T., Morgan, H. R., Andrews, J. C., Brewer, E. R. and Certel, S. J. (2017). Methyl-CpG binding domain proteins inhibit interspecies courtship and promote aggression in Drosophila. Sci. Rep., 7, 5420.

Hall, F. S. (1998). Social Deprivation of Neonatal, Adolescent, and Adult Rats Has Distinct Neurochemical and Behavioral Consequence. Crit. Rev. Neurobiol., 12, 129–62.

Hall, F. S., Wilkinson, L. S., Humby, T., Inglis, W., Kendall, D. A., Marsden, C. A. and Robbins, T. W. (1998). Isolation rearing in rats: Pre- and postsynaptic changes in striatal dopaminergic systems. Pharmacol. Biochem. Behav., 59, 859–872.

Hartigan, J. A. and Wong, M. A. (1979). Algorithm AS 136 A K-Means Clustering Algorithm. J. R. Stat. Soc. Ser. C (Applied Stat., 28, 100–108.

Hempel, C. M., Sugino, K. and Nelson, S. B. (2007). A manual method for the purification of fluorescently labeled neurons from the mammalian brain. Nat. Protoc., 2, 2924–2929.

Hendricks, J. C., Finn, S. M., Panckeri, K. A., Chavkin, J., Williams, J. A., Sehgal, A. and Pack, A. I. (2000). Rest in Drosophila Is a Sleep-like State. Neuron, 25, 129–138.

Henry, G. L., Davis, F. P., Picard, S. and Eddy, S. R. (2012). Cell type-specific genomics of Drosophila neurons. Nucleic Acids Res., 40, 9691–704.

Hu, Y., Flockhart, I., Vinayagam, A., Bergwitz, C., Berger, B., Perrimon, N. and Mohr, S. E. (2011). An integrative approach to ortholog prediction for disease-focused and other functional studies. BMC Bioinformatics, 12, 357.

Huang, D. W., Sherman, B. T. and Lempicki, R. A. (2009). Systematic and integrative analysis of large gene lists using DAVID bioinformatics resources. Nat. Protoc., 4, 44–57.

Johnson, A. A., Sarthi, J., Pirooznia, S. K., Reube, W. and Elefant, F. (2010). Increasing Tip60 HAT Levels Rescues Axonal Transport Defects and Associated Behavioral Phenotypes in a Drosophila Alzheimer’s Disease Model. J. Neurosci., 33, 7535–7547.

Jones, G. H., Hernandez, T. D., Kendall, D. A., Marsden, C. A. and Robbins, T. W. (1992). Dopaminergic and serotonergic function following isolation rearing in rats: Study of behavioural responses and postmortem and in vivo neurochemistry. Pharmacol. Biochem. Behav., 43, 17–35.

Jung, Y., Hsieh, L. S., Lee, A. M., Zhou, Z., Coman, D., Heath, C. J., Hyder, F., Mineur, Y. S., Yuan, Q., Goldman, D., et al. (2016). An epigenetic mechanism mediates developmental nicotine effects on neuronal structure and behavior. Nat. Neurosci., 19, 905–914.

Kaba, F., Lewis, A., Glowa-Kollisch, S., Hadler, J., Lee, D., Alper, H., Selling, D., MacDonald, R., Solimo, A., Parsons, A., et al. (2014). Solitary confinement and risk of self-harm among jail inmates. Am. J. Public Health, 104, 442–447.

Kadkhodaei, B., Ito, T., Joodmardi, E., Mattsson, B., Rouillard, C., Carta, M., Muramatsu, S.-I., Sumi-Ichinose, C., Nomura, T., Metzger, D., et al. (2009). Nurr1 Is Required for Maintenance of Maturing and Adult Midbrain Dopamine Neurons. J. Neurosci., 29, 15923–15932.

Kar, G., Kim, J. K., Kolodziejczyk, A. A., Natarajan, K. N., Triglia, E. T., Mifsud, B., Elderkin, S., Marioni, J. C., Pombo, A. and Teichmann, S. A. (2017). Flipping between Polycomb repressed and active transcriptional states introduces noise in gene expression. Nat. Commun., 8,.

Kayala, M. A. and Baldi, P. (2012). Cyber-T web server: Differential analysis of high-throughput data. Nucleic Acids Res., 40, 553–559.

Kharchenko, P. V, Alekseyenko, A. A., Schwartz, Y. B., Minoda, A., Riddle, N. C., Ernst, J., Sabo, P. J., Larschan, E., Gorchakov, A. A., Gu, T., et al. (2011). Comprehensive analysis of the chromatin landscape in Drosophila melanogaster. Nature, 471, 480–5.

Kim, J., Shin, S., Subramaniam, M., Bruinsma, E., Kim, T. D., Hawse, J. R., Spelsberg, T. C. and Janknecht, R. (2010). Histone demethylase JARID1B/KDM5B is a corepressor of TIEG1/KLF10. Biochem. Biophys. Res. Commun., 401, 412–416.

Koemans, T. S., Kleefstra, T., Chubak, M. C., Stone, M. H., F Reijnders, M. R., de Munnik, S., Willemsen, M. H., Fenckova, M., R M Stumpel, C. T., Bok, L. A., et al. (2017). Functional convergence of histone methyltransferases EHMT1 and KMT2C involved in intellectual disability and autism spectrum disorder. PLoS Genet., 1–24.

Kramer, J. M., Kochinke, K., Oortveld, M. a W., Marks, H., Kramer, D., de Jong, E. K., Asztalos, Z., Westwood, J. T., Stunnenberg, H. G., Sokolowski, M. B., et al. (2011). Epigenetic regulation of learning and memory by Drosophila EHMT/G9a. PLoS Biol., 9, e1000569.

Langmead, B. (2010). Aligning short sequencing reads with Bowtie. Curr. Protoc. Bioinforma., 32, 11.7.1–11.7.14.

Liao, Y., Smyth, G. K. and Shi, W. (2013). The Subread aligner: Fast, accurate and scalable read mapping by seed-and-vote. Nucleic Acids Res., 41, e108.

Liu, Q., Liu, S., Kodama, L., Driscoll, M. R. and Wu, M. N. (2012a). Two Dopaminergic Neurons Signal to the Dorsal Fan-Shaped Body to Promote Wakefulness in Drosophila. Curr. Biol., 22, 2114–2123.

Liu, C., Plaąais, P. Y., Yamagata, N., Pfeiffer, B. D., Aso, Y., Friedrich, A. B., Siwanowicz, I., Rubin, G. M., Preat, T. and Tanimoto, H. (2012b). A subset of dopamine neurons signals reward for odour memory in Drosophila. Nature, 488, 512–516.

Ma, J. and Weake, V. M. (2014). Affinity-based isolation of tagged nuclei from Drosophila tissues for gene expression analysis. J. Vis. Exp., 1–9.

Maze, I., Shen, L., Zhang, B., Garcia, B. a, Shao, N., Mitchell, A., Sun, H., Akbarian, S., Allis, C. D. and Nestler, E. J. (2014). Analytical tools and current challenges in the modern era of neuroepigenomics. Nat. Neurosci., 17, 1476–1490.

McGowan, P. O. and Szyf, M. (2010). The epigenetics of social adversity in early life: Implications for mental health outcomes. Neurobiol. Dis., 39, 66–72.

Mikkelsen, T. S., Ku, M., Jaffe, D. B., Issac, B., Lieberman, E., Giannoukos, G., Alvarez, P., Brockman, W., Kim, T.-K., Koche, R. P., et al. (2007). Genome-wide maps of chromatin state in pluripotent and lineage-committed cells. Nature, 448, 553–560.

Mo, A., Mukamel, E. A., Davis, F. P., Luo, C., Henry, G. L., Picard, S., Urich, M. A., Nery, J. R., Sejnowski, T. J., Lister, R., et al. (2015). Epigenomic Signatures of Neuronal Diversity in the Mammalian Brain. Neuron, 86, 1369–1384.

Mo, A., Luo, C., Davis, F. P., Mukamel, E. A., Henry, G. L., Nery, J. R., Urich, M. A., Picard, S., Lister, R., Eddy, S. R., et al. (2016). Epigenomic landscapes of retinal rods and cones. Elife, 5, 1–29.

Muñoz-Descalzo, S., Belacortu, Y. and Paricio, N. (2007). Identification and analysis of cabut orthologs in invertebrates and vertebrates. Dev. Genes Evol., 217, 289–298.

Nagoshi, E., Sugino, K., Kula, E., Okazaki, E., Tachibana, T., Nelson, S. and Rosbash, M. (2010). Dissecting differential gene expression within the circadian neuronal circuit of Drosophila. Nat. Neurosci., 13, 60–8.

Nitta, K. R., Jolma, A., Yin, Y., Morgunova, E., Kivioja, T., Akhtar, J., Hens, K., Toivonen, J., Deplancke, B., Furlong, E. E. M., et al. (2015). Conservation of transcription factor binding specificities across 600 million years of bilateria evolution. Elife, 4, e04837.

Niwa, M., Jaaro-Peled, H., Tankou, S., Seshadri, S., Hikida, T., Matsumoto, Y., Cascella, N. G., Kano, S., Ozaki, N., Nabeshima, T., et al. (2013). Adolescent Stress–Induced Epigenetic Control of Dopaminergic Neurons via Glucocorticoids. Science, 339, 335–340.

Niwa, M., Lee, R. S., Tanaka, T., Okada, K., Kano, S. I. and Sawa, A. (2016). A critical period of vulnerability to adolescent stress: Epigenetic mediators in mesocortical dopaminergic neurons. Hum. Mol. Genet., 25, 1370–1381.

Perkins, L. A., Holderbaum, L., Tao, R., Hu, Y., Sopko, R., McCall, K., Yang-Zhou, D., Flockhart, I., Binari, R., Shim, H. S., et al. (2015). The transgenic RNAi project at Harvard medical school: Resources and validation. Genetics, 201, 843–852.

Perry, S., Kiragasi, B., Dickman, D., Perry, S., Kiragasi, B., Dickman, D. and Ray, A. (2017). The Role of Histone Deacetylase 6 in Synaptic Plasticity and Memory. Cell Rep., 18, 1337–1345.

Pfeiffer, B. D., Truman, J. W. and Rubin, G. M. (2012). Using translational enhancers to increase transgene expression in Drosophila. Proc. Natl. Acad. Sci., 109, 6626–6631.

Pimentel, D., Donlea, J. M., Talbot, C. B., Song, S. M., Thurston, A. J. F. and Miesenböck, G. (2016). Operation of a homeostatic sleep switch. Nature, 536, 333–337.

Pusalkar, M., Suri, D., Kelkar, A., Bhattacharya, A., Galande, S. and Vaidya, V. A. (2015). Early stress evokes dysregulation of histone modifiers in the medial prefrontal cortex across the life span. Dev. Psychobiol.,

R Core Team (2014). R: A Language and Environment for Statistical Computing. R Found. Stat. Comput., Vienna,.

Reeves, R. and Tamburello, A. (2014). Single cells, segregated housing, and suicide in the New Jersey Department of Corrections. J. Am. Acad. Psychiatry Law, 42, 484–488.

Renthal, W., Kumar, A., Xiao, G., Wilkinson, M., Covington, H. E., Maze, I., Sikder, D., Robison, A. J., LaPlant, Q., Dietz, D. M., et al. (2009). Genome-wide analysis of chromatin regulation by cocaine reveals a role for sirtuins. Neuron, 62, 335–48.

Robinson, M. D., McCarthy, D. J. and Smyth, G. K. (2010). edgeR: a Bioconductor package for differential expression analysis of digital gene expression data. Bioinformatics, 26, 139–140.

Rodriguez, M. S., Dargemont, C. and Stutz, F. (2004). Nuclear export of RNA. Biol. Cell, 96, 639–655.

Sasagawa, T., Horii-Hayashi, N., Okuda, A., Hashimoto, T., Azuma, C. and Nishi, M. (2017). Long-term effects of maternal separation coupled with social isolation on reward seeking and changes in dopamine D1 receptor expression in the nucleus accumbens via DNA methylation in mice. Neurosci. Lett., 641, 33–39.

Schwartz, S., Truglio, M., Scott, M. J. and Fitzsimons, H. L. (2016). Long-Term Memory in Drosophila Is Influenced by Histone Deacetylase HDAC4 Interacting with SUMO-Conjugating Enzyme Ubc9. Genetics, 203, 1249–1264.

Shaw, P. J., Cirelli, C., Greenspan, R. J., Tononi, G., Campbell, S. S., Tobler, I., Zepelin, H., Rechtschaffen, A., Rechtschaffen, A., Tobler, I., et al. (2000). Correlates of sleep and waking in Drosophila melanogaster. Science, 287, 1834–7.

Shen, L., Shao, N.-Y., Liu, X., Maze, I., Feng, J. and Nestler, E. J. (2013). diffReps: Detecting Differential Chromatin Modification Sites from ChIP-seq Data with Biological Replicates. PLoS One, 8, e65598.

Shen, L., Shao, N., Liu, X. and Nestler, E. (2014). ngs.plot: Quick mining and visualization of next-generation sequencing data by integrating genomic databases. BMC Genomics, 15, 284.

Singh, A. S., Shah, A. and Brockmann, A. (2018). Honey bee foraging induces upregulation of early growth response protein 1, hormone receptor 38 and candidate downstream genes of the ecdysteroid signalling pathway. Insect Mol. Biol., 27, 90–98.

Sitaraman, D., Aso, Y., Rubin, G. M. and Nitabach, M. N. (2015). Control of Sleep by Dopaminergic Inputs to the Drosophila Mushroom Body. Front. Neural Circuits, 9, 1–8.

Siuda, D., Wu, Z., Chen, Y., Guo, L., Linke, M., Zechner, U., Xia, N., Reifenberg, G., Kleinert, H., Forstermann, U., et al. (2014). Social isolation-induced epigenetic changes in midbrain of adult mice. J. Physiol. Pharmacol., 65, 247–55.

Spain, M. M., Caruso, J. A., Swaminathan, A. and Pile, L. A. (2010). Drosophila SIN3 isoforms interact with distinct proteins and have unique biological functions. J. Biol. Chem., 285, 27457–27467.

Spittau, B., Wang, Z., Boinska, B. and Krieglstein, K. (2007). Functional domains of the TGF-β-inducible transcription factor Tieg3 and detection of two putative nuclear localization signals within the zinc finger DNA-binding domain. J. Cell. Biochem., 101, 712–722.

Steiner, F. A., Talbert, P. B., Kasinathan, S., Deal, R. B. and Henikoff, S. (2012). Cell-type-specific nuclei purification from whole animals for genome-wide expression and chromatin profiling. Genome Res., 22, 766–777.

Taniguchi, H. and Moore, A. W. (2014). Chromatin regulators in neurodevelopment and disease: Analysis of fly neural circuits provides insights: Networks of chromatin regulators and transcription factors underlie Drosophila neurogenesis and cognitive defects in intellectual disability and neuro. BioEssays, 36, 872–83.

Trapnell, C., Roberts, A., Goff, L., Pertea, G., Kim, D., Kelley, D. R., Pimentel, H., Salzberg, S. L., Rinn, J. L. and Pachter, L. (2012). Differential gene and transcript expression analysis of RNA-seq experiments with TopHat and Cufflinks. Nat. Protoc., 7, 562–78.

Ueno, T., Tomita, J., Tanimoto, H., Endo, K., Ito, K., Kume, S. and Kume, K. (2012). Identification of a dopamine pathway that regulates sleep and arousal in Drosophila. Nat. Neurosci., 15, 1516–1523.

Valzania, A., Catale, C., Viscomi, M. T., Puglisi-Allegra, S. and Carola, V. (2017). Histone deacetylase 5 modulates the effects of social adversity in early life on cocaine-induced behavior. Physiol. Behav., 171, 7–12.

van der Voet, M., Nijhof, B., Oortveld, M. A. W. and Schenck, A. (2014). Drosophila models of early onset cognitive disorders and their clinical applications. Neurosci. Biobehav. Rev., 46, 326–342.

Waddell, S. (2013). Reinforcement signalling in Drosophila; dopamine does it all after all. Curr. Opin. Neurobiol., 23, 324–329.

Wallace, D. L., Han, M.-H., Graham, D. L., Green, T. a, Vialou, V., Iñiguez, S. D., Cao, J.-L., Kirk, A., Chakravarty, S., Kumar, A., et al. (2009). CREB regulation of nucleus accumbens excitability mediates social isolation-induced behavioral deficits. Nat. Neurosci., 12, 200–9.

Wang, Y., Krishnan, H. R., Ghezzi, A., Yin, J. C. P. and Atkinson, N. S. (2007). Drug-induced epigenetic changes produce drug tolerance. PLoS Biol., 5, e265.

Weaver, I. C. G., Cervoni, N., Champagne, F. a, D’Alessio, A. C., Sharma, S., Seckl, J. R., Dymov, S., Szyf, M. and Meaney, M. J. (2004). Epigenetic programming by maternal behavior. Nat. Neurosci., 7, 847–54.

White, K. E., Humphrey, D. M. and Hirth, F. (2010). The dopaminergic system in the aging brain of Drosophila. Front. Neurosci., 4, 1–12.

Xu, S., Grullon, S., Ge, K. and Peng, W. (2014a). Spatial Clustering for Identification of ChIP-Enriched Regions (SICER) to Map Regions of Histone Methylation Patterns in Embryonic Stem Cells. Methods Mol. Biol., 1150, 97–111.

Xu, S., Wilf, R., Menon, T., Panikker, P., Sarthi, J. and Elefant, F. (2014b). Epigenetic control of learning and memory in drosophila by Tip60 HAT action. Genetics, 198, 1571–86.

Ye, Y., Li, M., Gu, L., Chen, X., Shi, J., Zhang, X. and Jiang, C. (2017). Chromatin remodeling during in vivo neural stem cells differentiating to neurons in early Drosophila embryos. Cell Death Differ., 24, 409–420.

Zang, C., Schones, D. E., Zeng, C., Cui, K., Zhao, K. and Peng, W. (2009). A clustering approach for identification of enriched domains from histone modification ChIP-Seq data. Bioinformatics, 25, 1952–1958.

Zetterström, R. H., Williams, R., Perlmann, T. and Olson, L. (1996). Cellular expression of the immediate early transcription factors Nurr1 and NGFI-B suggests a gene regulatory role in several brain regions including the nigrostriatal dopamine system. Mol. Brain Res., 41, 111–120.

Zhu, L. J., Christensen, R. G., Kazemian, M., Hull, C. J., Enuameh, M. S., Basciotta, M. D., Brasefield, J. A., Zhu, C., Asriyan, Y., Lapointe, D. S., et al. (2011). FlyFactorSurvey: A database of Drosophila transcription factor binding specificities determined using the bacterial one-hybrid system. Nucleic Acids Res., 39, D111–D117.

